# Endosomal cargo recycling mediated by Gpa1 and Phosphatidylinositol-3-Kinase is inhibited by glucose starvation

**DOI:** 10.1101/2021.04.02.438183

**Authors:** Kamilla ME. Laidlaw, Katherine M. Paine, Daniel D. Bisinski, Grant Calder, Karen Hogg, Sophia Ahmed, Sally James, Peter J. O’Toole, Chris MacDonald

**Affiliations:** York Biomedical Research Institute and Department of Biology, University of York, York, UK; Department of Biology & Chemistry, University of Osnabrück, Osnabrück, Germany; Bioscience Technology Facility, Department of Biology, University of York, UK

## Abstract

Cell surface protein trafficking is regulated in response to nutrient availability, with multiple pathways directing surface membrane proteins to the lysosome for degradation in response to suboptimal extracellular nutrients. Internalised protein and lipid cargoes recycle back to the surface efficiently in glucose replete conditions, but this trafficking is attenuated following glucose starvation. We find cells with either reduced or hyperactive phosphatidylinositol 3-kinase (PI3K) activity are defective for recycling. Furthermore, we find the yeast Gα subunit Gpa1, an endosomal PI3K effector, is required for surface recycling of cargoes. Following glucose starvation, mRNA and protein levels of a distinct Gα subunit Gpa2 are elevated following nuclear translocation of Mig1, which inhibits recycling of various cargoes. As Gpa1 and Gpa2 interact at the surface where Gpa2 concentrates during glucose starvation, we propose this disrupts PI3K activity required for recycling, potentially diverting Gpa1 to the surface and interfering with its endosomal role in recycling. In support of this model, glucose starvation and over-expression of Gpa2 alters PI3K endosomal phosphoinositide production. Glucose deprivation therefore triggers a survival mechanism to increase retention of surface cargoes in endosomes and promote their lysosomal degradation.

## INTRODUCTION

The surface localisation and activity of plasma membrane (PM) proteins can be regulated by the balance of action between endocytic trafficking pathways. Clathrin dependent and independent mechanisms internalise proteins from the PM, which then transit through different compartments *en route* to the lysosome for degradation. Various recycling mechanisms transport cargoes back to the surface, providing multiple regulatable steps to fine tune the surface protein environment in response to external conditions. Yeast has been a useful model organism to uncover conserved trafficking mechanisms of surface proteins. Upon internalisation from the yeast PM, surface cargoes can be sorted to the lysosome-like vacuole, in a process that involves cargo ubiquitination, mediated by E1-E2-E3 enzyme cascade in collaboration with competing trafficking adaptors (MacDonald *et al*., 2020; Sardana and Emr, 2021). Ubiquitinated cargoes are recognised at multivesicular bodies (MVBs) by the Endosomal Sorting Complex Require for Transport (ESCRT) apparatus, which also package cargo destined for degradation into intraluminal vesicles (Laidlaw and MacDonald, 2018). Proteins that are not targeted for degradation can recycle back to the surface, including a retrograde route that traffics material to the surface via the *trans*-Golgi network (TGN) using dedicated machineries that interact with recycled cargoes (Chen *et al*., 2019). Recycling in animal cells can also occur directly from early endosomes or indirectly, first traversing defined recycling endosomes (MacDonald and Piper, 2016). Endosomal organisation and recycling mechanisms in yeast are less clear (Ma and Burd, 2019a), but work using the yeast exocytic v-SNARE protein Snc1 revealed multiple endosomal transport steps regulate Snc1 trafficking back to the PM (Ma *et al*., 2017; Ma and Burd, 2019b; Best *et al*., 2020). Although retrograde recycling is perturbed by deubiquitination (Xu *et al*., 2017), recycling of some nutrient transporters is triggered by deubiquitination (MacDonald and Piper, 2017; Laidlaw *et al*., 2021). Genetic dissection of this latter pathway implies recycling is controlled at transcriptional and metabolic levels (MacDonald and Piper, 2017; Amoiradaki *et al*., 2021), suggesting early endocytic trafficking decisions contribute to the eventual downregulation of surface cargoes in response to nutritional cues.

During periods of nutritional stress, multiple mechanisms involving surface cargoes and endocytic pathways are modulated to promote proliferation, particularly in cancer cells (Selwan *et al*., 2016; Finicle *et al*., 2018). Recycling internalised cargoes back to the surface can promote anabolic processes. During starvation, when some such processes are not required, reduced recycling can route surface cargoes to the lysosomal (vacuolar in yeast) degradation pathway instead, to promote catabolism. Conceptually, these pathways can be modulated to drive growth/proliferation appropriate to extracellular nutrient availability. One such example has been elucidated in yeast responding to nitrogen starvation, where increased trafficking to the vacuole is achieved through the amino acid sensing TORC1 complex, which activates Rsp5-adaptors via the Npr1 kinase and promotes cargo ubiquitination and degradation (MacGurn *et al*., 2011). The Rag GTPases integrate with nutrient sensing and TORC1 activity via the EGO complex (Binda *et al*., 2009; Bonfils *et al*., 2012; Péli-Gulli *et al*., 2015), but in addition regulate recycling via endosomally-localised Ltv1 in response to extracellular Leucine (MacDonald and Piper, 2017). Glucose starvation may also trigger a similar dual regulation of recycling and degradation pathways in yeast, where many metabolic pathways are evolutionarily conserved (Santangelo, 2006). Although most transcriptional changes elucidated in response to suboptimal glucose availability involve alternative carbon pathways, we recently revealed a response that increases trafficking from the surface to the lysosome in response to glucose starvation (Laidlaw *et al*., 2020). Furthermore, recycling back to the surface is reduced when cells are exposed to media lacking sugar (Lang *et al*., 2014), but the molecular players involved in this response are not fully established.

In high glucose levels, Snf1 (yeast AMP-activated protein kinase, AMPK (Hong *et al*., 2003)) is inactivated primarily through its dephosphorylation by Reg1-Glc7 (Tu and Carlson, 1994; Sanz *et al*., 2000). Whilst Snf1 can influence endosomal trafficking during glucose changes in yeast independently (O’Donnell *et al*., 2015), one key downstream consequence of glucose starvation involves Snf1 activation and regulation of the Mig1 transcriptional repressor (Johnston *et al*., 1994; Treitel *et al*., 1998). Mig1 is a zinc finger transcription factor that mediates glucose repression in yeast cells (Nehlin and Ronne, 1990; Lundin *et al*., 1994) and in response activates alternative metabolic pathways (Schüller, 2003). In addition to this, Mig1 influences membrane trafficking machinery through its repression of clathrin adaptor genes *YAP1801* and *YAP1802* under glucose replete conditions that stabilise endocytosis levels (Laidlaw *et al*., 2020). Furthermore, the reduced recycling following glucose starvation (Lang *et al*., 2014) can be bypassed by enforcing cargo ubiquitination in the endomembrane system upstream of MVB sorting (Buelto *et al*., 2020). This suggests that both the ubiquitin-mediated degradation pathway (MacDonald and Piper, 2016), and the counteracting recycling pathway induced following cargo deubiquitination (MacDonald *et al*., 2012a, 2015a) are both regulated in response to glucose starvation to collectively reduce surface activity and increased vacuolar degradation.

A genetic screen for recycling machinery identified several candidates for glucose-mediated control, including the G-protein coupled receptor (GPCR) Gpr1 and downstream GTPase Ras2 (MacDonald and Piper, 2017). The Gα subunit Gpa2, which is activated Gpr1, initiates cAMP signalling in response to glucose through recruiting Ras-GTP to the PM (Colombo *et al*., 1998; Broggi *et al*., 2013). This cAMP signalling cascade in response to glucose leads to changes in a variety of different targets through protein kinase A (PKA) (Thevelein, 1994; Kraakman *et al*., 1999). The screen for recycling machinery also implicated a distinct Gα subunit, Gpa1 in recycling (MacDonald and Piper, 2017). Gpa1 is the Gα subunit of a heterotrimeric G protein complex also comprised of Gβ and Gγ subunits Ste4p and Ste18p, that functions in the pheromone response pathway (Dietzel and Kurjan, 1987; Miyajima *et al*., 1987; Whiteway *et al*., 1989). Just as Gpa2 cannot functionally couple with distinct mating GPCRs (Blumer and Thorner, 1990) it is thought that the distinct the Gα subunit Gpa1 does not regulate Gpr1 directly. In addition to playing a role in pheromone response at the PM, Gpa1 interacts with the yeast phosphatidylinositol 3-kinase (Vps15 and Vps34) at endosomes (Slessareva *et al*., 2006; Heenan *et al*., 2009). The phosphorylation status of a variety of different phosphoinositide (PI) species regulates membrane trafficking pathways (Camilli *et al*., 1996). Phosphoinositide 3-kinases (PI3Ks) are therefore key regulatory proteins known to control a variety of different membrane trafficking steps (Lindmo and Stenmark, 2006). Vps34, in complex with Vps15, was first identified in yeast as being required for the post-Golgi trafficking of biosynthetic enzymes to the vacuole (Herman and Emr, 1990; Schu *et al*., 1993; Stack *et al*., 1993) and generation of phosphatidylinositol 3-phosphates (PtdIns3P) through Vps34 activity is required for efficient retrograde recycling from the endosome to the late-Golgi (Burda *et al*., 2002).

In this study, we use both lipid and protein recycling reporters to show that glucose starvation inhibits endosomal recycling of cargo back to the surface. We show the Gα subunit Gpa1 and PI3K, and their functional association, are required for efficient recycling in glucose replete conditions. During glucose starvation, we document Mig1-dependent elevation of Gpa2, which concentrates at the PM where it physically interacts with Gpa1. Increased levels of Gpa2 observed in glucose starved cells impairs production of PtdIns3P and results in recycling defects of a wide range of cargoes. We propose a role for Gpa2 as a glucose responsive recycling inhibitor, potentially by commandeering Gpa1 from endosomal PI3K, and ultimately serving to increases endosomal retention and vacuolar degradation of surface cargoes, as a survival response to glucose starvation.

## RESULTS

### Glucose starvation inhibits surface recycling

Previous work has shown that in response to glucose starvation, the yeast AP180 clathrin adaptors are transcriptionally upregulated with a concurrent increase in endocytosis (Laidlaw *et al*., 2020). Under basal conditions in media lacking methionine, the methionine transporter Mup1 tagged with GFP localises to the plasma membrane (PM), but much of this signal is redistributed to endosomes and the vacuole upon plasmid over-expression of mCherry-tagged AP180s: Yap1801-Cherry or Yap1802-mCherry (**Figure 1A**). In contrast, a recycling reporter based on the fusion of the G-protein couple receptor (GPCR) Ste3 tagged with GFP and the catalytic domain of a deubiquitinating enzyme (Stringer and Piper, 2011; MacDonald and Piper, 2017), showed no increase in endosomal localisation following over-expression of yeast AP180s (**Figure 1B**). This correlates with observations that Ste3-GFP-DUb does not accumulate when endocytosis rates are elevated by the addition of **a**-factor or at elevated temperature (MacDonald and Piper, 2017). We conclude that although different manipulations increase the rate of endocytosis, the deubiquitination-driven recycling of Ste3-GFP-DUb predominates to maintain an exclusive steady state localisation at the PM. Having established that Ste3-GFP-DUb primarily reports on cell surface recycling, we used this reporter to test whether glucose starvation impacts recycling specifically. Treating the cells in media completely lacking sugar, or by substituting glucose for the alternative carbon course raffinose, results in an accumulation of Ste3-GFP-DUb in intracellular endosomes (**Figure 1C**). A distinct recycling assay, which measures recycling by efflux of endocytosed fluorescent FM4-64 (Wiederkehr *et al*., 2000), has previously shown carbon source removal inhibits recycling (Lang *et al*., 2014). Similarly, we find shifting cells to media lacking sugar or supplemented with raffinose also results in robust inhibition of recycling (**Figure 1D**). Therefore, although internalisation from the PM increases following glucose starvation (Laidlaw *et al*., 2020), two dedicated recycling reporters show glucose starvation also triggers a reduction in protein and lipid traffic from endosomes back to the PM. We propose increased internalisation and decreased recycling during glucose starvation cooperate to drive vacuolar degradation of cargoes *en masse* in response to nutritional stress. We set out to explain this recycling response at a molecular level. As both glucose removal and raffinose exhibit defects in recycling, we use raffinose substitution to deprive cells of glucose in downstream experiments, as it cannot be readily metabolised (Fuente and Sols, 1962) and therefore in the time-frame we perform experiments provides a glucose starvation condition.

**Figure 1:**
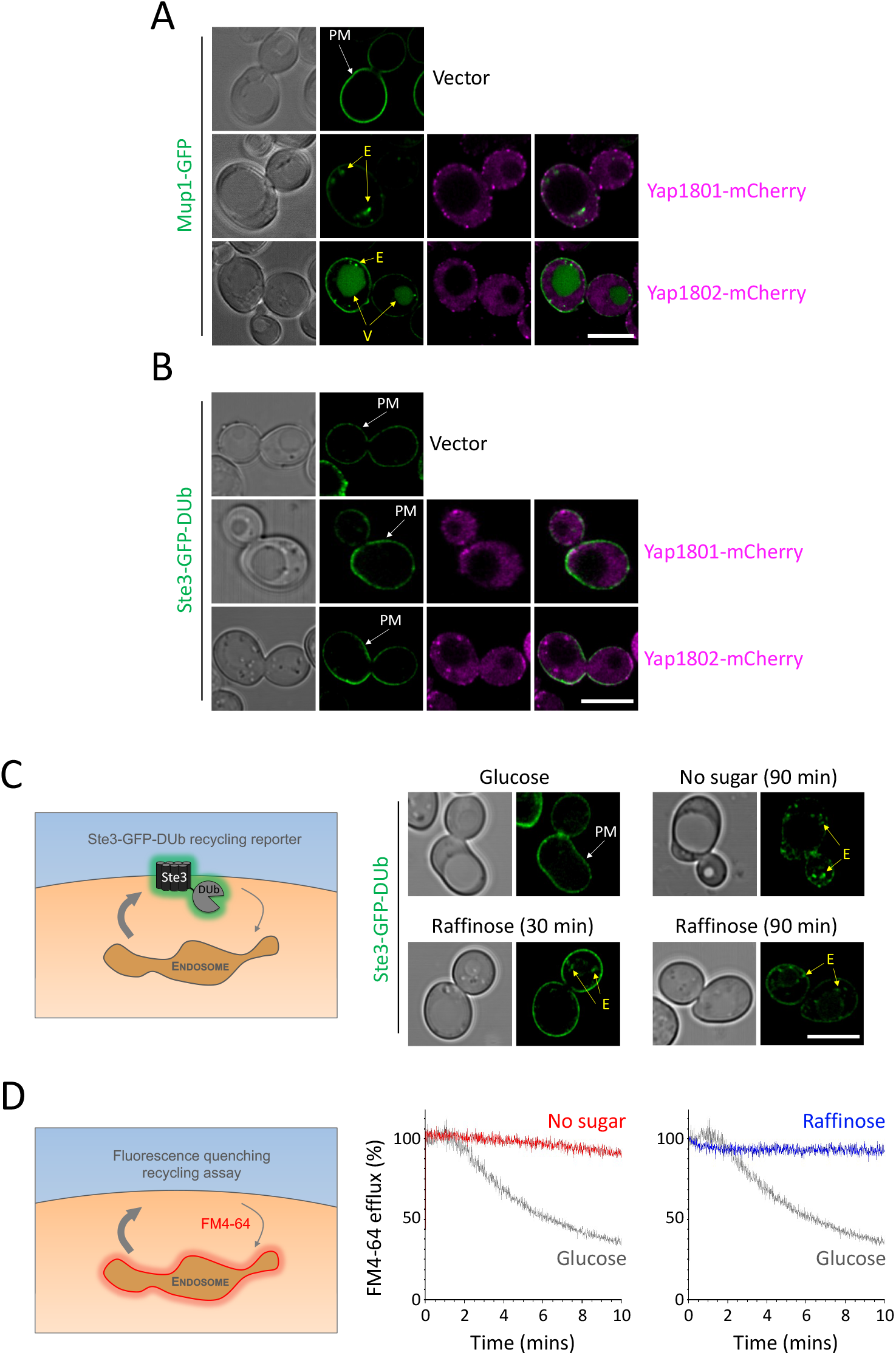
Glucose starvation specifically inhibits recycling. **A)** Confocal imaging of cells expressing Mup1-GFP under control of its endogenous promoter co-expressed with Yap1801-mCherry or Yap1802-mCherry expressed from the *CUP1* promoter through addition of 50 µM copper chloride for 2 hours. A vector control exposed to copper was also included. **B)** Cells stably integrated with Ste3-GFP-DUb were exposed to 2 hours copper chloride to induce expression of mCherry tagged versions of Yap1801 and Yap1802, before confocal microscopy. **C)** Cells expressing Ste3-GFP-DUb from the *STE3* promoter were grown under indicated media conditions prior to confocal microscopy. **D)** Wild-type cells loaded with rich media containing 40 µM FM4-64 for 8-minutes before washing and dye efflux measured over time by flow cytometry. Control cells grown in glucose media were compared with 30-minute prior incubation with media lacking any carbon source (red) or media supplemented with raffinose (blue). White arrows indicate exclusive plasma membrane (PM) signal and yellow arrows indicate endosome (E) or vacuole (V) localisations. Scale bar, 5 µm.

### Gpa1-PI3K is required for surface recycling

A previous genetic screen for novel recycling machinery, based on the mis-localisation of Ste3-GFP-DUb, (MacDonald and Piper, 2017) identified 89 candidate factors as required for recycling (**Figure 2A**). To identify any proteins from this list that might inhibit recycling during glucose starvation (**Figure 1**), a network analysis of all 89 proteins was performed based on both physical and functional associations to reveal a small cluster, containing the glucose regulated receptor Gpr1, and associated factors Ras2 and Gpa1 (**Figure 2B**). We stably integrated the Ste3-GFP-DUb reporter into strains lacking each factor (*gpa1Δ, ras2Δ*, and *gpr1Δ*) and confirmed they exhibit defects in recycling (**Figure 2C, 2D**). Although deletion of these genes results in a pronounced mis-localisation phenotype, we also performed flow cytometry experiments to confirm that total levels of Ste3-GFP-DUb were not elevated to account for additional signal in intracellular compartments (**Figure 2E**). Although all three candidates are known to localise and function at the PM, we focussed on the Gα subunit Gpa1 for this study, as it has also been shown to localise to endosomes and activate PI3K (Slessareva *et al*., 2006), which is involved in various endomembrane trafficking events (Lindmo and Stenmark, 2006; Reidick *et al*., 2017). We first considered the G-protein subunit Gpa1 might specifically perturb the Ste-GFP-DUb reporter, as it based on the G-protein coupled receptor Ste3. However, we found general recycling of lipids, as assessed by FM4-64 efflux, was also defective in *gpa1Δ* mutants (**Figure 2F**). Furthermore, trafficking of endogenous cargoes to the PM was also perturbed in *gpa1Δ* cells. We found the methionine transporter Mup1 tagged with GFP, which localises exclusively to the PM in wild-type cells, is shifted to endosomes and the vacuole in *gpa1Δ* mutants (**Figure 2G**). Similarly, the PM signal of the uracil transporter Fur4, tagged with mNeonGreen (mNG) (Paine *et al*., 2021), or Ste3 tagged with GFP (but lacking a deubiquitinating enzyme fusion), was sorted to the vacuole in *gpa1Δ* cells. These defects in lipid and protein trafficking are consistent with a role for Gpa1 in surface recycling.

**Figure 2:**
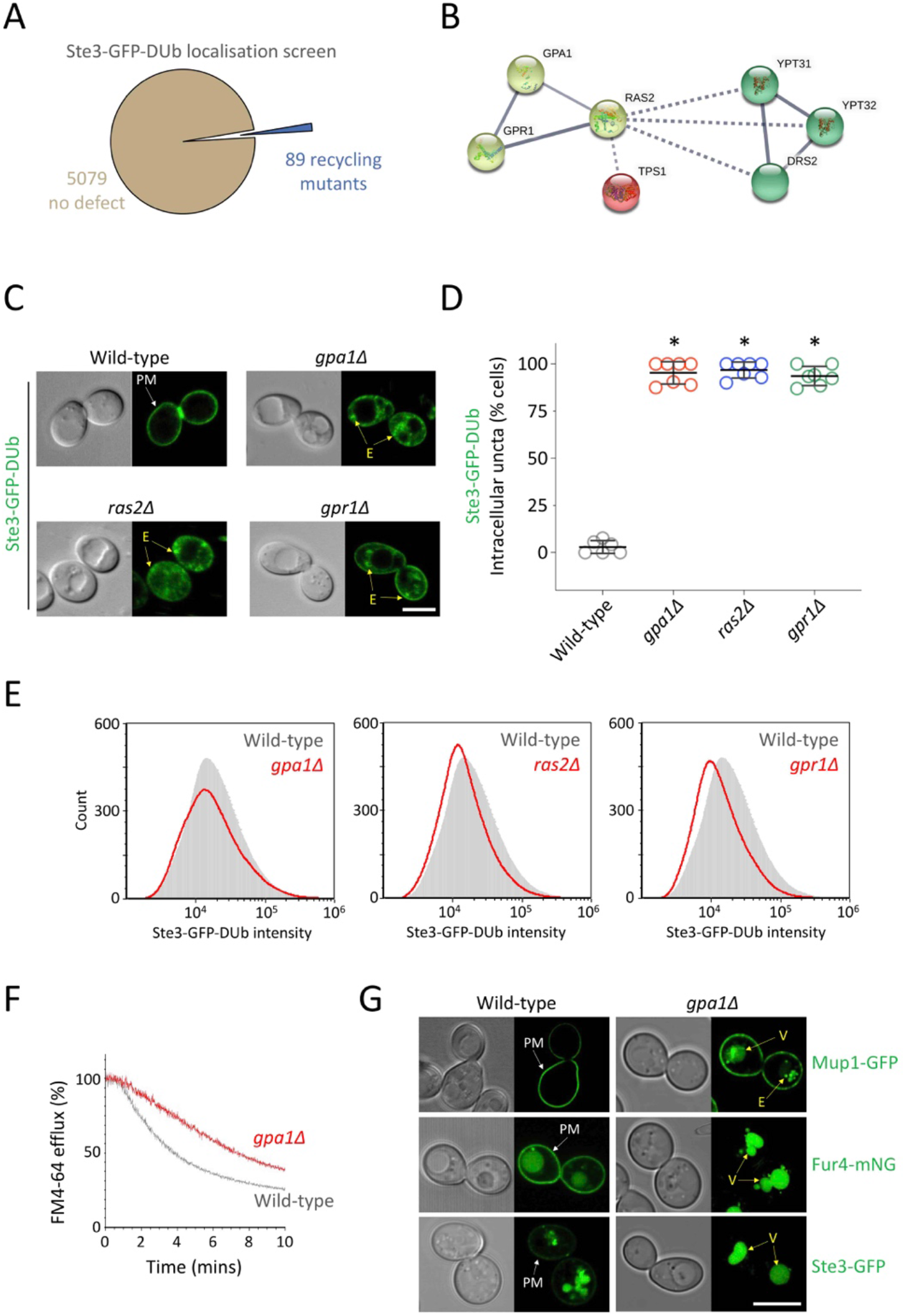
Gpa1 is required for protein and lipid recycling to the surface. **A)** Pie chart representing gene deletions that have no impact on Ste3-GFP-DUb recycling (brown), or those that accumulate the reporter in endosomes (blue). **B)** String pathway analysis was performed on all 89 recycling factor candidates (minimum interaction score = high confidence, 0.700) before application of a k-means clustering algorithm to define 9 groups. A small cluster connected by solid lines, representing strongly supported functional and physical associations (yellow) is shown alongside associations with distinct clusters (red and green) shown by broken lines. **C)** Quantification of Ste3-GFP-DUb intracellular localisations calculated as an average of population (n=3) from WT = 90; *gpa1Δ* = 75; *ras2Δ* = 94; and *gpr1Δ* = 70 cells. **D)** Ste3-GFP-DUb localisation recorded in indicated cells by confocal microscopy. **E)** Flow cytometry was used to measure Ste3-GFP-DUb fluorescence from approximately 75,000 cells of each indicated mutant (red) and compared with expression in wild-type cells (grey overlay). **F)** Wild-type (grey) or *gpa1Δ* (red) cells were loaded with FM4-64 for 8-minutes before dye efflux was assessed by flow cytometry over 10-minutes. **G)** Wild-type and *gpa1Δ* cells expressing Mup1-GFP, Fur4-mNG or Ste3-GFP were grown to log phase before confocal microscopy to determine localisation. White arrows (exclusive PM) and yellow arrows (vacuole; V) are indicated. Scale bar, 5 µm.

Constitutive signalling in haploid *gpa1Δ* mutants is lethal (Miyajima *et al*., 1987) so we first confirmed *GPA1* deletion in both our Mat**a** and Mat***α*** backgrounds by genotyping (**Figure 3A**) and genome sequencing (**Figure 3B**). We surveyed all genetic mutations identified from both *gpa1Δ* mutants and wild-type cells and found most variants were far from ORFs, so unlikely to affect expression, or missense variants unlikely to alter function. Gene ontology of mutated genes revealed a premature stop codon in the *STE11* gene of Mat**a** *gpa1Δ* cells (**Figure 3C, 3D**), which would perturb downstream signalling and suppress lethality (Nakayama *et al*., 1988). Explanation of suppression in Mat***α*** mutants is less clear but might be due to a substitution in the downstream mitogen activated protein (MAP) kinase kinase (MAPKK) *MKK1*. This alone, or in combination with a parental strain point mutation in *SST2*, which encodes a negative regulator of Gpa1-signalling (Dohlman *et al*., 1996; Apanovitch *et al*., 1998), might explain viability of *gpa1Δ* mutants.

**Figure 3:**
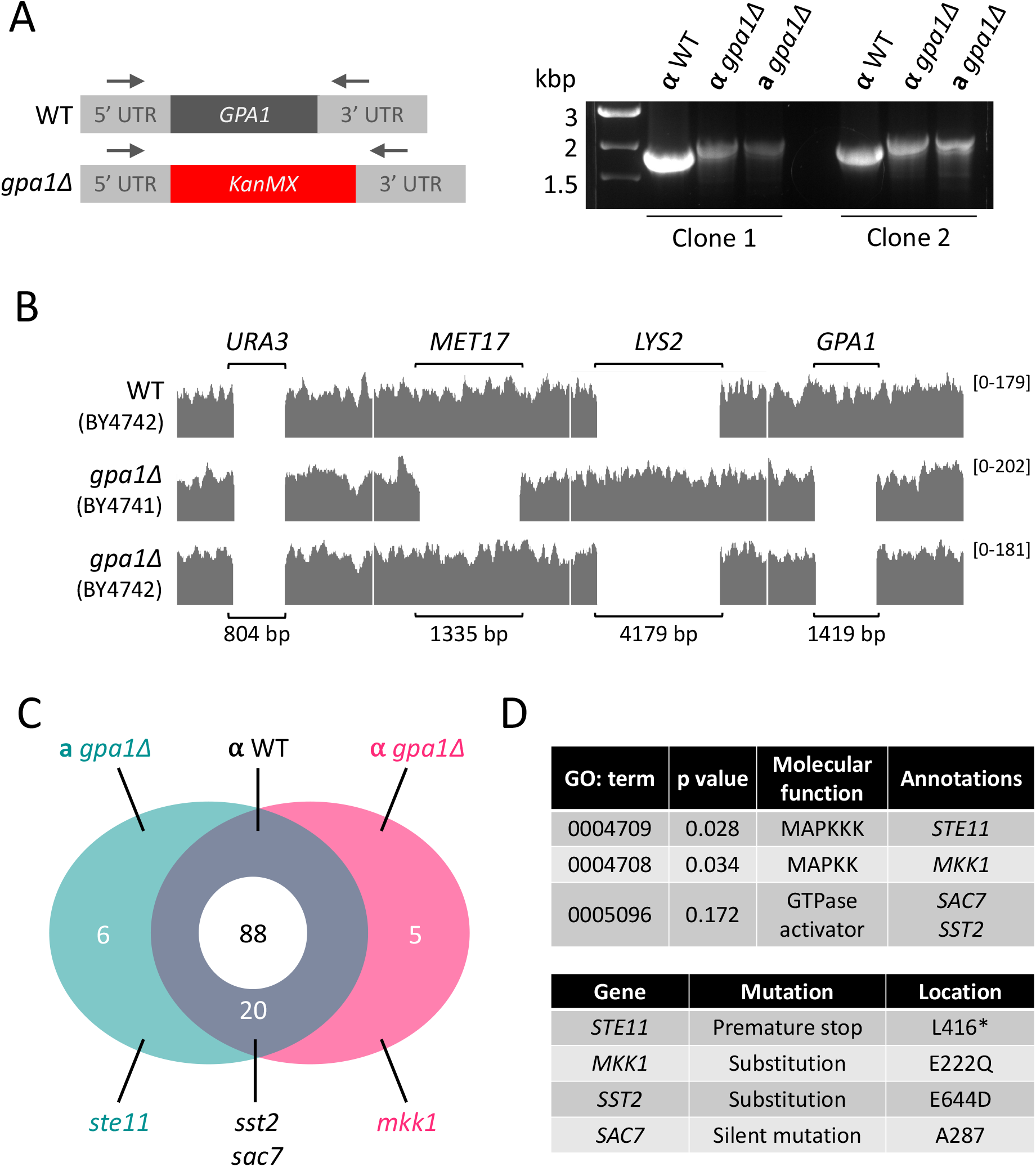
Genetic validation of viable *gpa1Δ* mutant yeast strains. **A)** Schematic depicting region of the genome that was PCR amplified using oligos in 5’ and 3’ UTRs of *GPA1* (Left). gDNA was isolated from Mat**α** (wild-type and *gpa1Δ*) and Mat**a** *gpa1Δ* cells and used for PCR confirmation of the *KanMX* deletion cassette in *gpa1Δ* mutants (right). **B)** Genome sequencing was performed on cells described in **(A)** with read-depth visualised in Integrative Genomics Viewer (IGV) software for specific loci shown: *URA3*, deleted in both BY-parental strains; *MET17* deleted in BY4741; *LYS2* deleted in BY4742; and *GPA1*, deleted in *gpa1Δ* cells in both mating type. **C)** Venn diagram depicting the overlapping gene mutations related to Gpa1 found in sequenced strains. **D)** List of enriched annotations from gene ontology (GO) analysis related to mating and Gpa1-signalling (upper) and specific mutation details of identified mutations (lower).

To test if these Gpa1 effects are mediated through PI3K, we analysed recycling in cells lacking PI3K subunits (*vps15Δ* and *vps34Δ*), which both exhibit morphological defects of the vacuolar / endolysosomal system (**Figure 4A**). Vacuole morphology following FM4-64 staining was difficult to resolve by conventional confocal microscopy but Airyscan microscopy revealed layers of small vacuolar-like structures in both *vps15Δ* and *vps34Δ* cells. Both *vps15Δ* and *vps34Δ* mutants were confirmed to have a growth defect at 30°C (**Figure 4B**) before we revealed that *vps15Δ* and *vps34Δ* cells are severely defective in their ability to recycle internalised FM4-64 dye (**Figure 4C**). To further corroborate this model, we employed a hyperactive version of Vps34, termed Vps34^EDC^ (harbouring R283E, A287D, Y501C point mutations) that was recently used to show PtdIns3P production is rate limiting for some, but not all, membrane trafficking pathways (Steinfeld *et al*., 2021). We found that unlike deletion of PI3K, increased expression of Vps34^WT^ or Vps34^EDC^ had no effect on growth at 30°C (**Figure 4B**). However, expressing hyperactive Vps34^EDC^ was sufficient to perturb efficient recycling of FM4-64 (**Figure 4D**) and Ste3-GFP-DUb (**Figure 4E**), suggesting elevated PtdIns3P production deregulates recycling, but to a lesser degree than in cells lacking PI3K activity. To test if a functional connection between Gpa1 and PI3K is required for efficient recycling, we created an R1261A mutation in endogenous Vps15 (**Figure 4F, 4G**), which disrupts interaction with Gpa1 (Heenan et al., 2009), and found this was sufficient to inhibit efficient FM4-64 recycling (**Figure 4H**). This supports the notion that yeast PI3K in collaboration with Gpa1 is responsible for recycling from endosomes to the surface.

**Figure 4:**
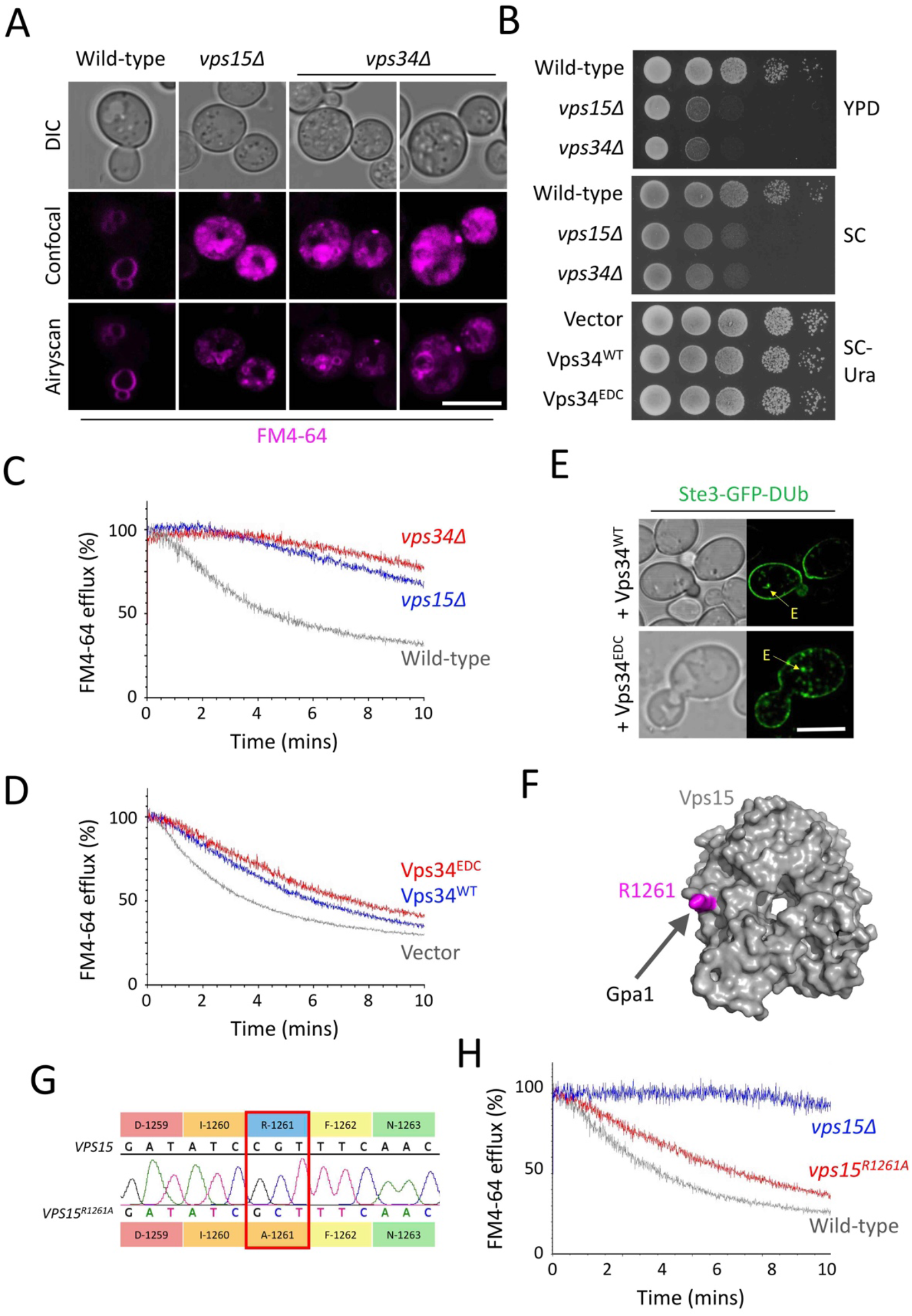
Defined PI3-Kinase activity is required for efficient surface recycling. **A)** Confocal and Airyscan imaging of wild-type, *vps15Δ* and *vps34Δ* cells first labelled with 10 µM FM4-64 for 30 minutes, followed by washing and a 1-hour chase in SC minimal media. **B)** Indicated cells were grown to log phase, equivalent volumes harvested and a 1 in 10 serial dilution spotted onto YPD and synthetic complete (SC) media and incubated at 30°C. **C**,**D**,**H)** Indicated strains and transformants were loaded with 40 µM FM4-64 in rich media for 8-minutes, before 3x 5 minute washes and FM4-64 dye efflux measured over time by flow cytometry and plotted by the % of the initial 10 s fluorescence. **E)** Wild-type cells co-expressing Ste3-GFP-DUb with either Vps34^WT^ or Vps34^EDC^ were imaged using confocal microscopy. **F)** Surface model of Vps15 WD-repeat domain (grey) with R1261 shown (magenta). **G)** Sequencing results of the *vps15*^*R1261A*^ allele confirmed by Sanger sequencing of PCR product generate by PCR. Scale bar, 5 µm.

### Reg1 regulates recycling at transcriptional level

Gpa1 localises to the PM and endomembrane compartments (Slessareva *et al*., 2006; Dixit *et al*., 2014). Fluorescently tagged Gpa1, which functionally complements *gpa1Δ* cells (**Figure S1**), colocalises with the late endosome marker Vps4, and to a lesser degree, the TGN marker Sec7 (**Figure 5A**). We found there was a subtle defect in Ste3-GFP-DUb recycling upon over-expression of Gpa1-mCherry (**Figure 5B**), suggesting both PI3K and Gpa1 levels need to be finely tuned for efficient recycling back to the surface. Intracellular Ste3-GFP-DUb colocalised with Gpa1-mCherry, in both wild-type and recycling defective *rcy1Δ* mutants, suggesting a trafficking block in endosomes from which recycling occurs. Expressing a constitutively active version (Gpa1^Q323L^-mCherry) caused more severe defects in Ste3-GFP-DUb (**Figure 5C, 5F**), again with endosomal recycling reporter colocalise Gpa1. Intriguingly, the Gpa1^Q323L^ screen that revealed Gpa1 couples with PI3K during PtdIns3P production (Slessareva *et al*., 2006) also demonstrated the glucose-related phosphatase *REG1* is functionally associated Gpa1 (**Figure 5D**). To explore whether the recycling defects associated with Gpa1-PI3K are related to glucose mediated recycling, we tested recycling in *reg1Δ* mutants and found both Ste3-GFP-DUb (**Figure 5E, 5F**) and FM4-64 (**Figure 5F**) do not efficiently recycle. No obvious recycling defects were observed in cells lacking the distinct *SNF3* glucose sensing component. We hypothesised the connection between Glucose / Reg1 and Gpa1 / PI3K allowed cargo recycling from endosomes to the surface to be regulated in response to available glucose.

**Figure 5:**
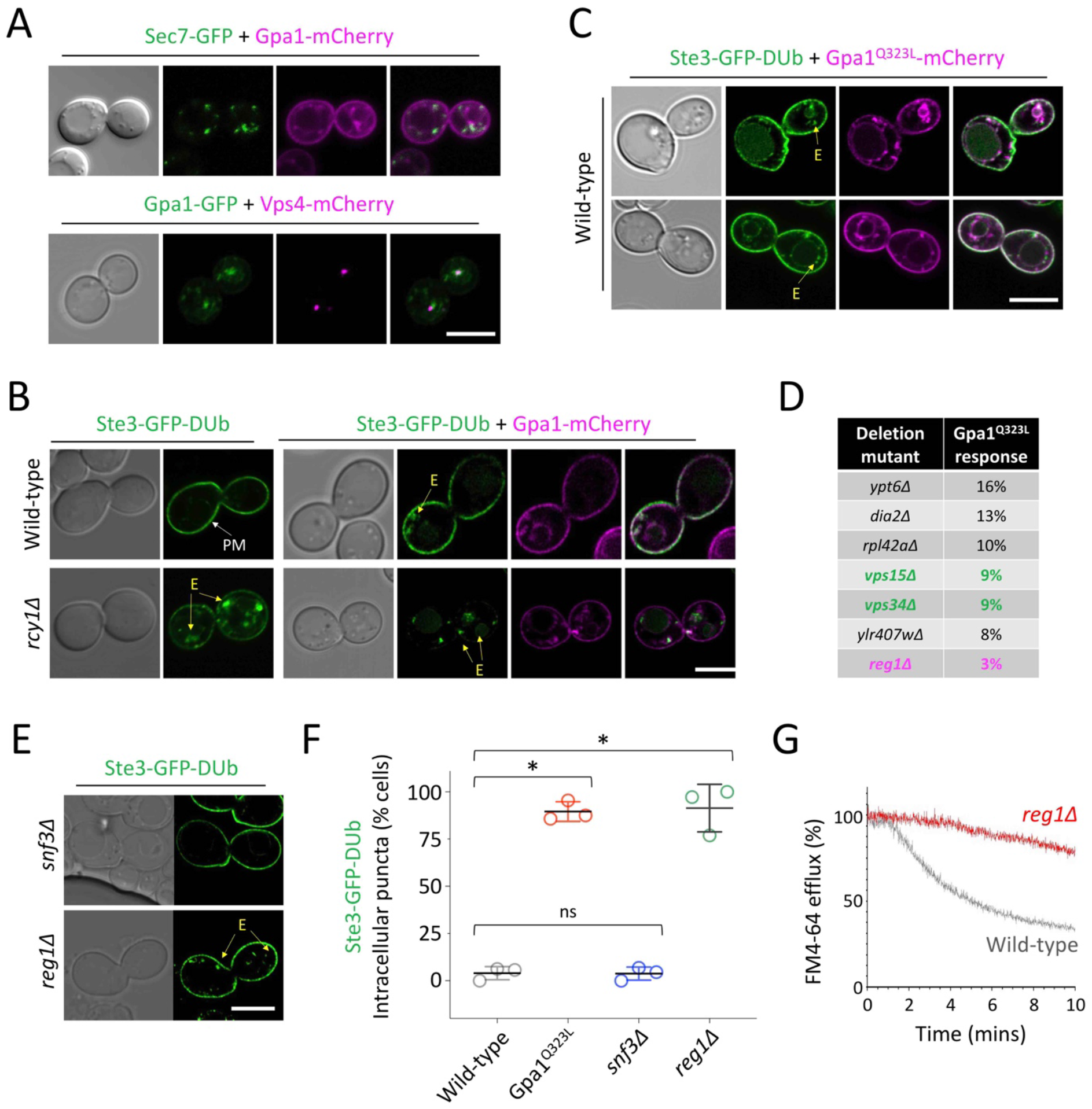
Reg1 is a proposed upstream Gpa1 regulator in recycling. **A)** Confocal microscopy of wild-type cells co-expressing Sec7-GFP and Gpa1-mCherry (top) and Gpa1-GFP and Vps4-mCherry (bottom). **B)** Wild-type and *rcy1*Δ cells expressing Ste3-GFP-DUb with and without Gpa1-mCherry were imaged by confocal microscopy. **C)** Ste3-GFP-DUb localisation in wild-type cells co-expressing Gpa1^Q323L^-mCherry was assessed by fluorescence microscopy. **D)** Top scoring mutants from a Gpa1^Q323L^ mating response screen (Slessareva *et al*., 2006) are shown, with PI3K mutants (green) and *reg1*Δ (magenta) highlighted. **E)** Confocal microscopy of *snf3Δ* and *reg1Δ* cells stably expressing Ste3-GFP-DUb. **F)** Quantification of Ste3-GFP-DUb intracellular localisations calculated as an average of population (n=3) from WT = 68; Gpa1^Q323L^ = 51; *snf3Δ* = 83; and *reg1Δ* = 73 cells. **G)** Wild-type (grey) and *reg1Δ* (red) cells were incubated with rich media containing 40 µM FM4-64 for 8-minutes before washing and FM4-64 dye efflux measured over time by flow cytometry and plotted by the % of the initial 10 s fluorescence. Scale bar, 5 µm.

Reg1, alongside the essential phosphatase Glc7 and the yeast AMPK family member Snf1, participate in a signal transduction pathway that controls transcription in response to glucose (Tu and Carlson, 1994, 1995; Ludin *et al*., 1998; Sanz *et al*., 2000). Snf1 regulates the Mig1 transcriptional repressor in response to glucose levels to repress various metabolic genes (Johnston *et al*., 1994; Vallier and Carlson, 1994; Treitel *et al*., 1998). We considered a model whereby this signalling mechanism maintained high levels of surface cargoes in glucose replete conditions by repression of a recycling inhibitor that acts on Gpa1-PI3K (**Figure 6A**). We went on to identify one candidate inhibitor (discussed below): the Gα subunit Gpa2, which physically and genetically interacts with Gpa1 (Xue et al., 1998; Ho et al., 2002) that we propose fulfils the function of Gpa1-recycling inhibitor. This model would predict that deletion of *MIG1* and *MIG2* repressors (Lutfiyya *et al*., 1998; Westholm *et al*., 2008), would phenocopy the defective recycling observed followed by glucose starvation or in either *reg1Δ* or *gpa1Δ* cells. As a glucose responsive inhibitor, transcript levels of *GPA2* would increase Gpa2 protein levels following glucose starvation, which would consequently inhibit Gpa1-recycling (**Figure 6B**). This model would predict that elevated levels of PM localised Gpa2 interacting with Gpa1 would hamper the finely tuned production of PtdIns3P via Gpa1-PI3K required for efficient surface recycling.

**Figure 6:**
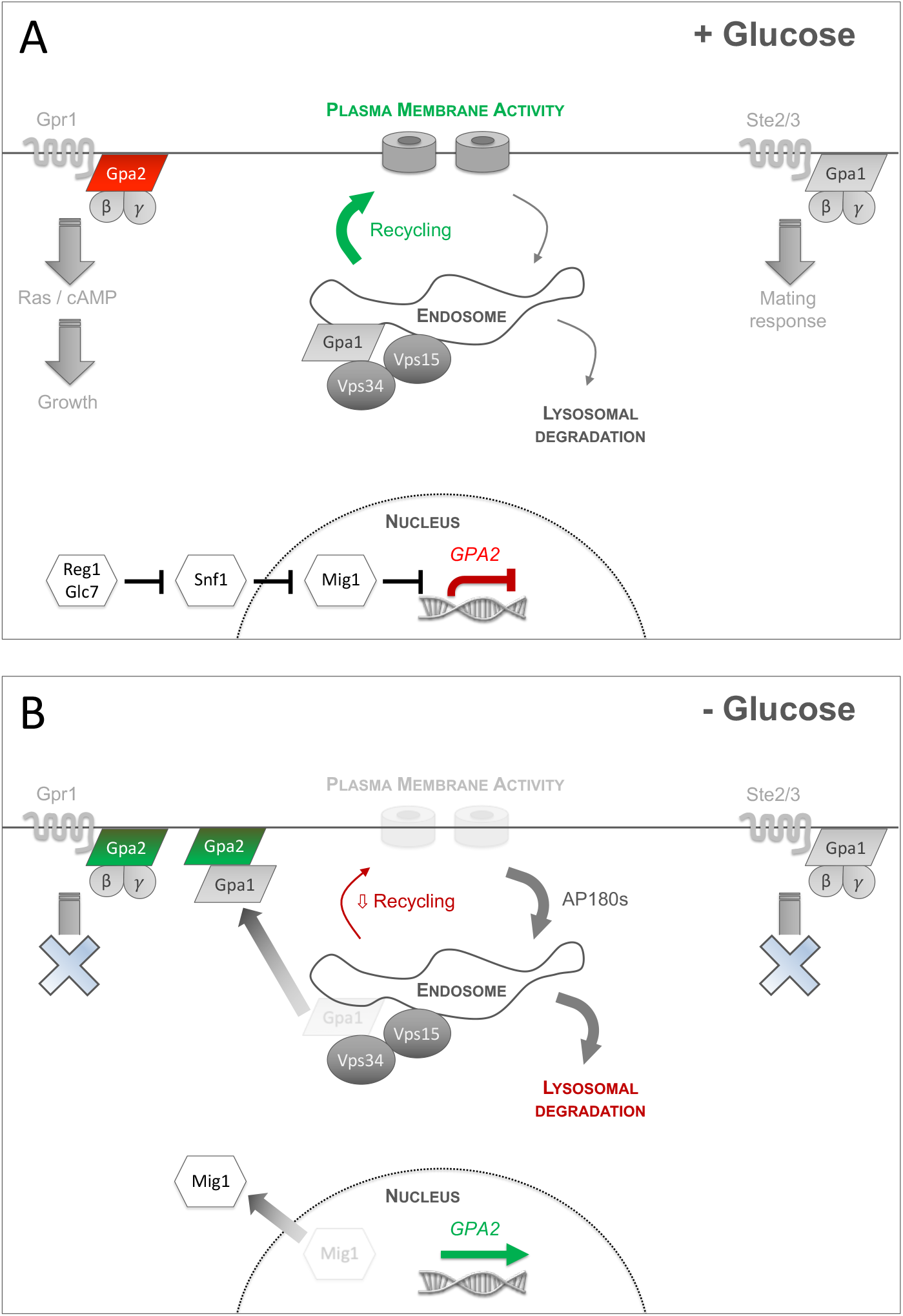
Model for glucose mediated control of cargo recycling. **A)** In glucose rich conditions, metabolism of the cell is maintained to promote growth, in part through the cAMP synthesis pathway sensed through the GPCR Gpr1 and heterotrimeric G protein alpha subunit, Gpa2. In glucose conditions, we propose *GPA2* expression is suppressed via the established glucose repression pathway involving Glc7/Reg1 > Snf1 > Mig1. The pheromone signalling and mating pathway, controlled through haploid specific GPCRs Ste2/Ste3 also function at the PM via the G protein alpha subunit, Gpa1. Gpa1 also has a function at endosomes, with PI3Kinase subunits Vps15/Vps34 producing PtdIns3P. Surface recycling of PM cargoes is efficient in glucose replete conditions, via an active and efficient recycling pathway from endosomes back to the surface. **B)** Glucose starvation triggers several metabolic changes, including growth arrest sensed via Gpr1. This is accompanied by a reduction in mating response, as the Ste2/3 receptors are downregulated. The Glc7 > Snf1 pathway results in dephosphorylation and translocation of nuclear repressor Mig1. In consequence to this, the yeastAP180 clathrin adaptors are transcriptionally upregulated and induce higher levels of internalisation from the PM. Concomitantly, levels of *GPA2* are increased, which we propose acts as an inhibitor of Gpa1-mediated recycling, potentially through sequestering more Gpa1 at the PM and therefore decamping from PI3kinase at the endosome and disrupting the lipid organisation required to efficiently promote recycling.

### Gpa2 inhibits surface recycling

We find recycling defects of *gpa1Δ* cells are phenocopied in *reg1Δ* mutants (**Figure 2C, 2E, 5E, 5F**). We reasoned if the functional role of Reg1 in recycling is through its capacity to modulate transcription via the Reg1/Glc7 > Snf1 > Mig1/2 pathway, then recycling defects of *reg1Δ* cells would also be phenocopied in *mig1Δ mig2Δ* mutant cells. Alternatively, if recycling is efficient in *mig1Δ mig2Δ* cells, a direct role between Reg1, and its substrate Gpa1 (Clement *et al*., 2013), might best explain the data. Mig1 is a glucose sensitive transcriptional repressor that rapidly translocate from the nucleus when cells are shifted to raffinose media (**Figure 7A, 7B, S2**). Many glucose repressed genes are transcriptionally upregulated upon glucose starvation or in *mig1Δ mig2Δ* cells lacking the repressors (Westholm *et al*., 2008). As *mig1Δ mig2Δ* mutants exhibit perturbed Ste3-GFP-DUb recycling (**Figure 7C**), we assume recycling defects in *reg1Δ* mutants are explained via a transcriptional response, as discussed (**Figure 6**). To identify genes transcriptionally controlled by Reg1>Mig1 that inhibit Gpa1-mediated recycling, we cross-referenced a list of genes predicted to be repressed by Mig1 (Wollman *et al*., 2017) with the physical interactome of Gpa1 to reveal only one candidate, Gpa2 (**Figure 7D**). To test whether *GPA2* was a *bona fide* target gene for Mig1 repression controlled by glucose we performed qPCR experiments. Firstly, we revealed that *GPA1* transcript levels are unchanged in response to Mig1-repression or available glucose. In contrast, the proposed Gpa1-inhibitor *GPA2* is transcriptionally upregulated ∼2.0 ± 0.3 fold in *mig1Δ mig2Δ* cells and ∼6.4 ± 0.7 fold upon a shift to raffinose for 1-hour (**Figure 7E**). This transcriptional profile is consistent with over-expression of a Gpa1-recycling-inhibitor. Similarly, deletion of the proposed inhibitor exhibits no defects in Ste3-GFP-DUb recycling (**Figure 7F**), suggesting only increasing levels of Gpa2 inhibits recycling. We did find *gpa1Δ gpa2Δ* cells are defective for Ste3-GFP-DUb recycling, supporting the idea that whilst Gpa1-PI3K is required for recycling, Gpa2 is a downstream regulator. The Ste3-GFP-DUb recycling defects triggered by deletion of *MIG1* and *MIG2* can be attributed to the induced levels of Gpa2 in these cells, as additionally deleting *GPA2* in a *mig1Δ mig2Δ* background suppresses recycling defects (**Figure 7G**). We found cells expressing Gpa2-GFP localised almost exclusively to the PM, where Gpa1-mCherry partially co-localises, in addition to its endosomal localisation. However, this endosomal population shifts to primarily PM in *rcy1Δ* recycling mutants (**Figure S3**).

**Figure 7:**
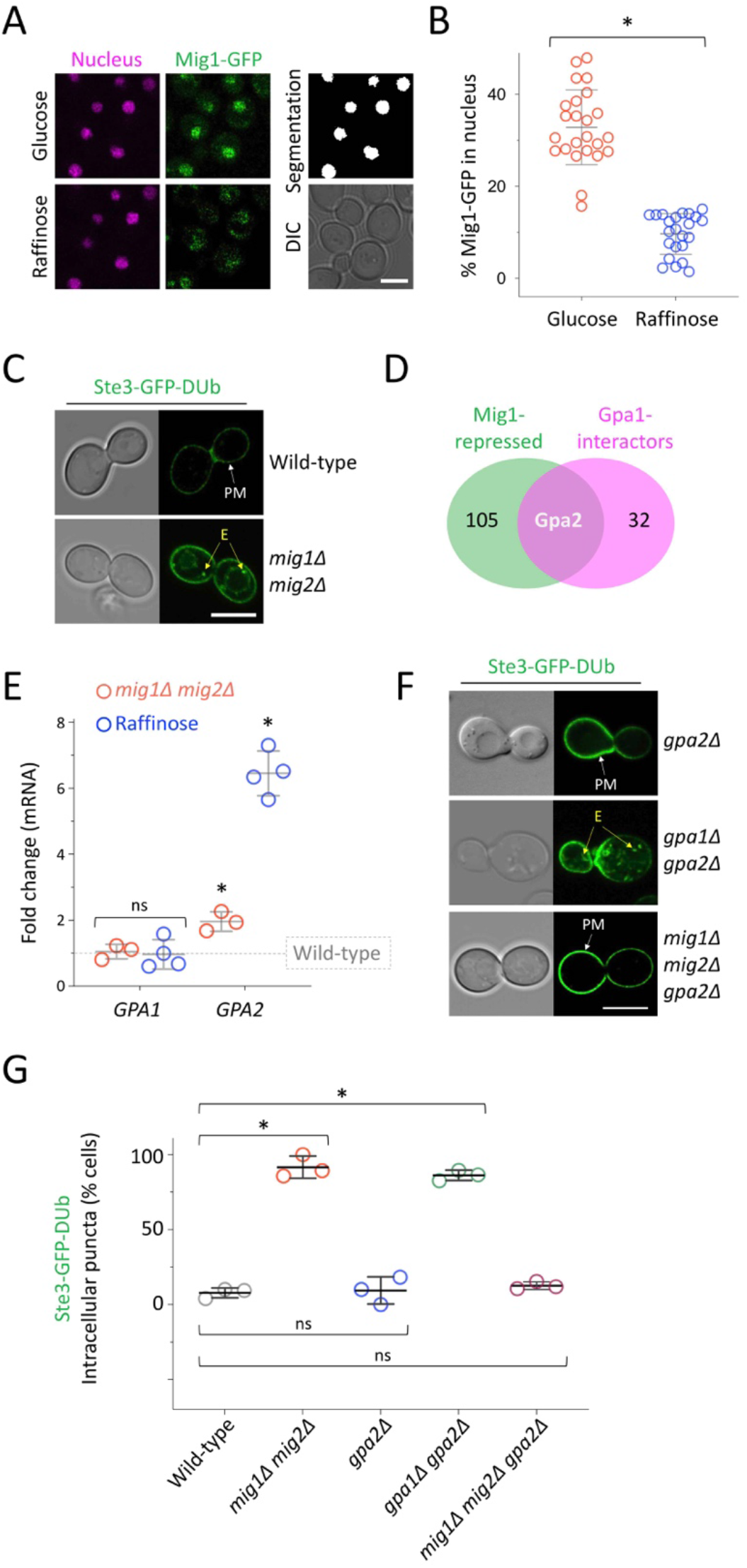
Glucose and Mig1-controlled expression of recycling inhibitor *GPA2*. **A)** Confocal microscopy of wild-type cells co-expressing Mig1-GFP (green) and Nrd1-mCherry as a nuclear marker (magenta) under glucose replete conditions. The same cells were imaged 5-minutes later following a raffinose exchange performed with microfluidics. **B)** The percentage of nuclear Mig1-GFP signal under glucose and raffinose conditions was quantified (see methods). * indicates unpaired Holm–Sidak *t*-test (p < 0.0001). **C)** Confocal microscopy of wild-type and *mig1Δmig2Δ* cells expressing Ste3-GFP-DUb. **D)** Venn diagram of Mig1-repressed candidates (green) and proteins that physically interact with Gpa1 (magenta). **E)** RT-qPCR was used to measure transcript levels of indicated genes, compared to *ACT1*, in wild-type versus *mig1Δ mig2Δ* (red) and wild-type cells grown in glucose versus 60-minutes raffinose media (blue). Unpaired Holm–Sidak *t*-tests showed *GPA1* levels were not significantly (ns) altered in either experiment (p = 0.748 and 0.907, respectively) but levels of *GPA2* increased in both *mig1Δ mig2Δ* (p = <0.001) and raffinose (p = < 0.00001). **F)** Confocal microscopy of indicated mutants expressing Ste3-GFP-DUb. **G)** Quantification of Ste3-GFP-DUb intracellular localisations calculated as an average of population (n=3) from WT = 77; *mig1Δ mig2Δ* = 64; *gpa2Δ* = 69; *gpa1Δ gpa2Δ* = 79; and *mig1Δ mig2Δ gpa2Δ* = 87 cells. Scale bar, 5 µm.

We next confirmed this Mig1-based transcriptional response results in an increase in Gpa2-GFP protein levels by immunoblot following 2-hours raffinose treatment (**Figure 8A, 8B**). By assessing maximum intensity projects from 3D Airyscan imaging of cells expressing Gpa2-GFP, we also found a pool of cytosolic Gpa2-GFP in glucose grown cells redistributes to the PM following raffinose treatment (**Figure 8C, 8D**). Similar analysis of 3D images focussing only on the top or bottom of cells also revealed punctate PM accumulations of Gpa2-GFP in raffinose grown cells (**Figure 8E, 8F**), but it is unclear if these have functional significance or are indirectly due to the influx of higher protein levels and greater surface/cytoplasm ratios. Our model predicts that elevated levels of Gpa2 would inhibit recycling, which was confirmed by over-expressing plasmid borne fluorescently tagged versions of Gpa2, which we confirmed are functional (**Figure S4**). We find over-expressed Gpa2-mCherry triggered accumulation of Ste3-GFP-DUb in intracellular compartments (**Figure 9A, 9B**). Similarly, cells over-expressing Gpa2-GFP have reduced efflux of FM4-64 from the recycling pathway, when compared to control cells expressing the methionine transporter Mup1-GFP (**Figure 9C, S5**). In addition to these recycling specific cargoes, we also used the Gpa2-mCherry over-expression system to reveal prominent defects in recycling of the PM localised GFP tagged transporters Mup1 and Can1, which shift to endosomes and the vacuole (**Figure 9D**). Gpa2 over-expression induced a reduction of Ste3-GFP from the PM, however we also note that vacuolar sorting, which is typically evident in wild-type cells at steady state, is blocked following Gpa2-mCherry overexpression, and cargo accumulates in prevacuolar structures. We assume overexpression of the Gα subunit Gpa2 perturbs trafficking of Ste3 in a manner distinct from general Gpa1-PI3K lipid-mediated trafficking. As Ste3 is a GPCR, excess Gpa2 might force it to adopt a conformation at the PM that is not conducive to MVB sorting.

**Figure 8:**
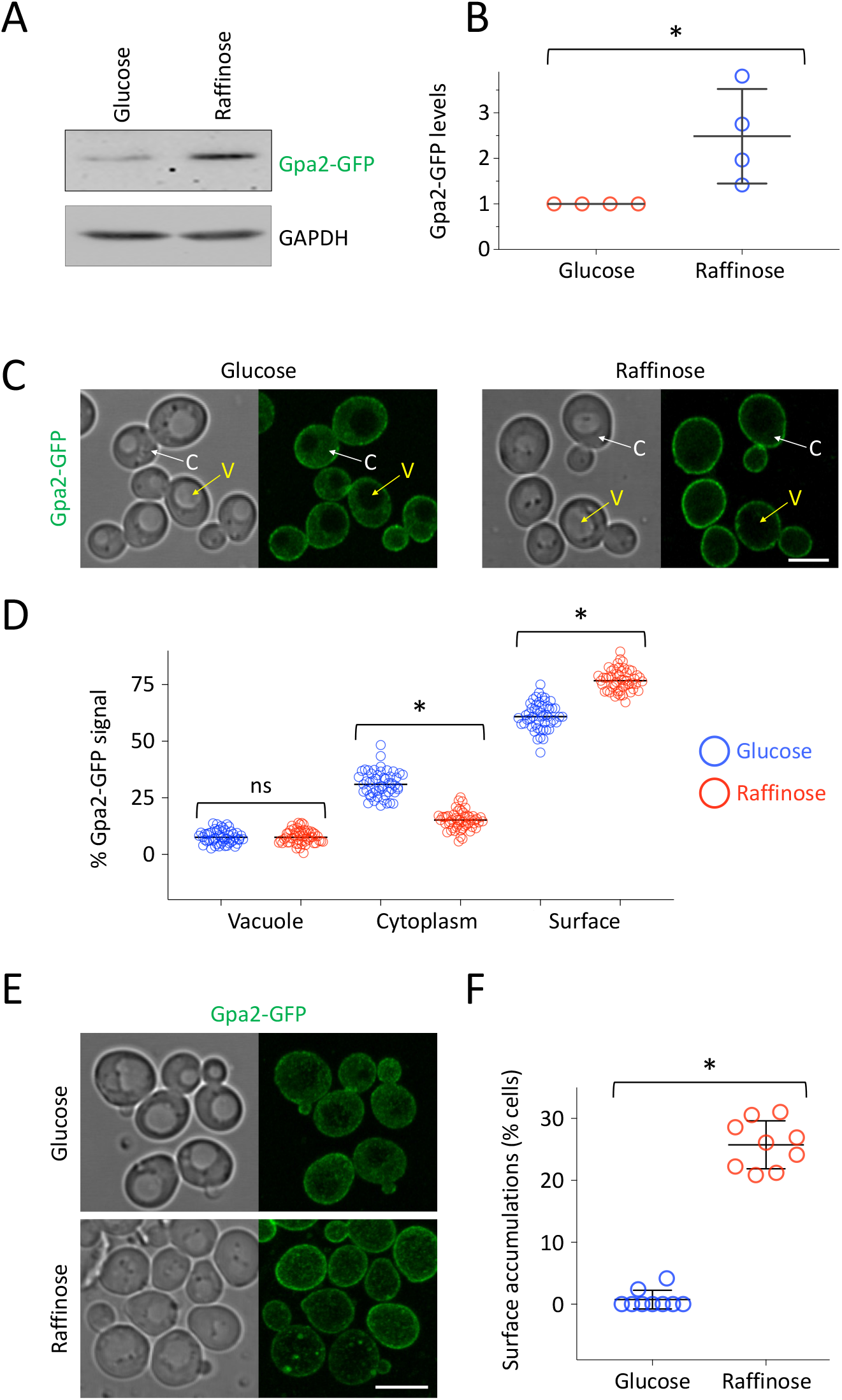
Glucose starvation induced expression and surface concentration of Gpa2-GFP. **A)** Wild-type cells expressing GFP tagged Gpa2 were grown in glucose and 90 minutes raffinose conditions before lysates were made for immunoblot analysis probing with **α**-GFP and **α**-GAPDH antibodies. **B)** Gpa2-GFP expression levels in glucose and 90 minutes raffinose treatment were quantified from immunoblots (A) using imageJ and normalised to GAPDH signal. * indicates unpaired *t-*test (*p* = 0.0287). **C)** Wild-type cells expressing endogenously expressed Gpa2-GFP were imaged by Airyscan in glucose (left) and following 2 hours raffinose treatment (right) were presented as max-intensity projections from 4 z-stack slices from centre focussed cells Arrows showing vacuole (V; yellow) and cytoplasm (C; white) are shown. Mean intensity measurements for vacuole, cytoplasm and surface from cells in **(C)** were performed. **E)** Max-intensity projections of 3D confocal images of 4 z-stack slices covering the top-focussed area of the cell. **F)** Cells exhibiting surface puncta of Gpa2-GFP were quantified as a percentage of total population. * indicates unpaired *t-*test (p < 0.0001). Scale bar, 5 µm.

**Figure 9:**
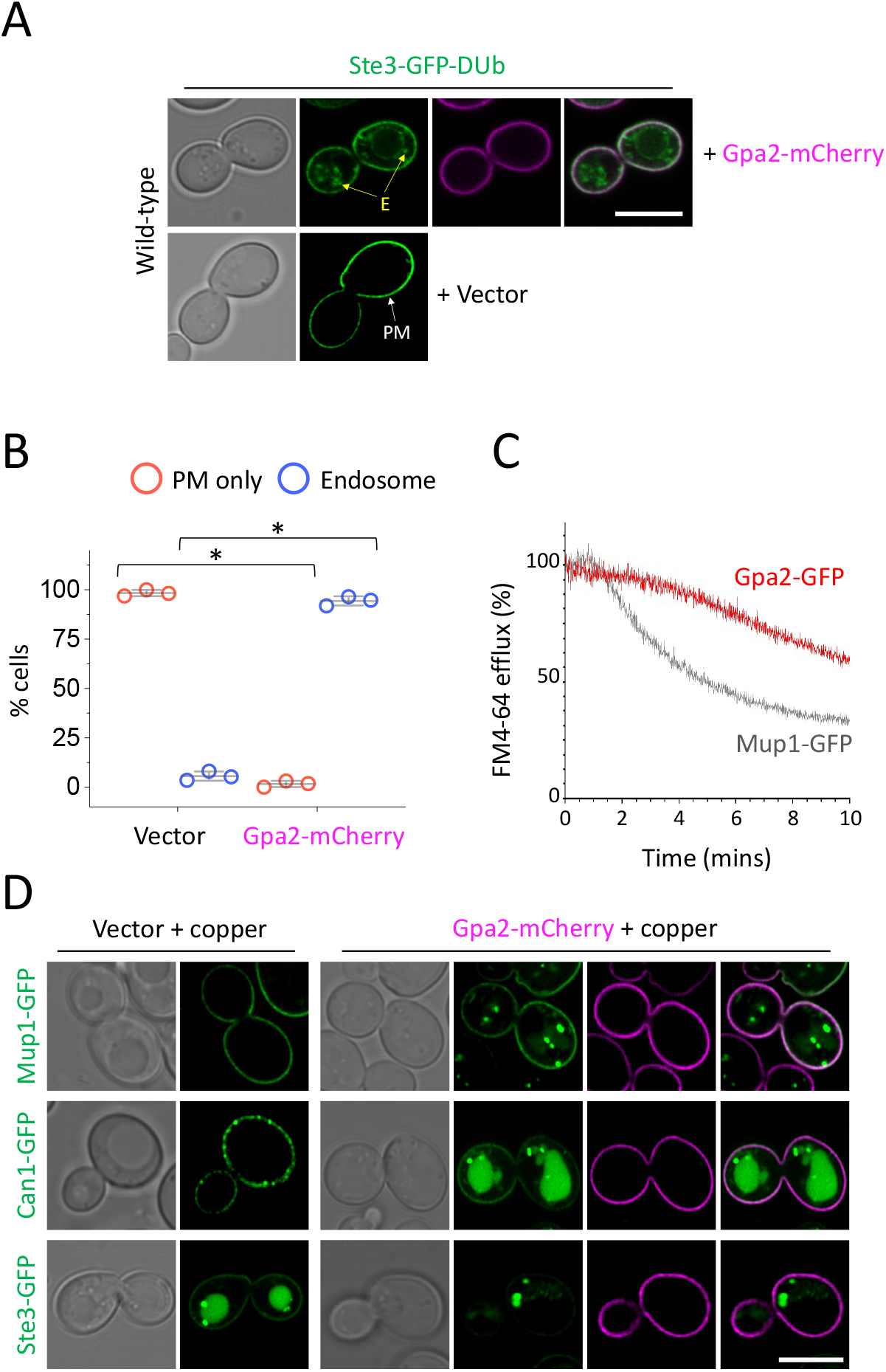
Over-expression of Gpa1 perturbs surface recycling. **A)** Confocal microscopy of wild-type cells expressing Ste3-GFP-DUb with and without Gpa2-mCherry. **B)** Localisation of Ste3-GFP-DUb in wild-type cells transformed with vector or Gpa2-mCherry at mid-log phase was quantified from >35 cells (n = 3). Unpaired Students *t*-tests were performed, with asterisk (*) indicating p < 0.000001. **C)** FM4-64 efflux assay was measured from wild-type cells expressing either Gpa2-GFP or Mup1-GFP. Cells were loaded with rich media containing 40 µM FM4-64 for 8-minutes before washing and dye efflux measured over time by flow cytometry and expressed a % of the initial 10 s fluorescence. **D)** Wild-type cells expressing the copper inducible Gpa2-mCherry construct, using 50 µM copper chloride, and co-expressing either Mup1-GFP, Can1-GFP or Ste3-GFP were imaged using confocal microscopy. Scale bar, 5 µm.

To explore whether Gpa2 could function at endosomes directly we performed time-lapse microscopy of cells expressing Gpa2-GFP and only very rarely observed intracellular puncta, however these did not colocalise with Gpa1-mCherry (**Figure 10A**). We did find strong accumulations of Gpa2-GFP within subdomains of the PM, which are distinct from Mup1-GFP localisation and partially colocalise with brief pulses of the endocytic dye FM4-64 (**Figure 10B**). In order to assess Gpa2-GFP localisation across the entire cell, we optimised fast yet gentle Apotome Structured Illumination Microscopy (SIM) imaging to capture fluorescence across the entire cell volume, with 42 distinct z-stack slices in both colour channels, captured in only 4.3 seconds. Initial experiments were calibrated using the vacuolar cargo Cos5 (MacDonald *et al*., 2015a). Cos5-GFP accumulates in the vacuole of wild-type cells but concentrates in class E endosomes in *vps25Δ* mutant cells (**Figure S6**), so we used a dual tagged Cos5-GFP-mCherry to optimise processing and noise correction (**Figure 10C**). 4D Apotome SIM microscopy experiments show Gpa2-GFP displays a continuous network distribution pattern across the PM (**Figure 10D**) reminiscent of the localisation of the Gpa2 associated protein Ras2 (Spira *et al*., 2012) but no co-localisation with mCherry tagged markers for the trans-Golgi network (TGN) and the multivesicular body (MVB) (**Figures 10E, S7**). As these efforts did not provide strong evidence for Gpa2 localising anywhere other than the surface, we propose the function of higher levels of Gpa2 at the PM during glucose starvation could simply be to appropriate more Gpa1, thereby depriving PI3K of Gpa1 and reducing its capacity to mediate efficient recycling. In support of this idea, we reveal a significant Förster Resonance Energy Transfer (FRET) signal between Gpa2-GFP and Gpa1-mCherry at the surface (**Figure 10F, 10G, S8**), but no indication of FRET between Gpa1 and Sec7 or in unbleached controls. Finally, we show that PI3K production of PtdIns3P is impaired in raffinose treated cells, indicated by the mis-localisation of the PX-domain protein Snx41 and the FYVE-domain protein Pib1(**Figure 11A, 11B**), which both bind endosomal membranes rich in PtdIns3P produced by PI3K (Burd and Emr, 1998; Shin *et al*., 2001; Yu and Lemmon, 2001; Hettema *et al*., 2003). These mis-localisation effects were phenocopied in glucose grown cells over-expressing Gpa2 from a plasmid.

**Figure 10:**
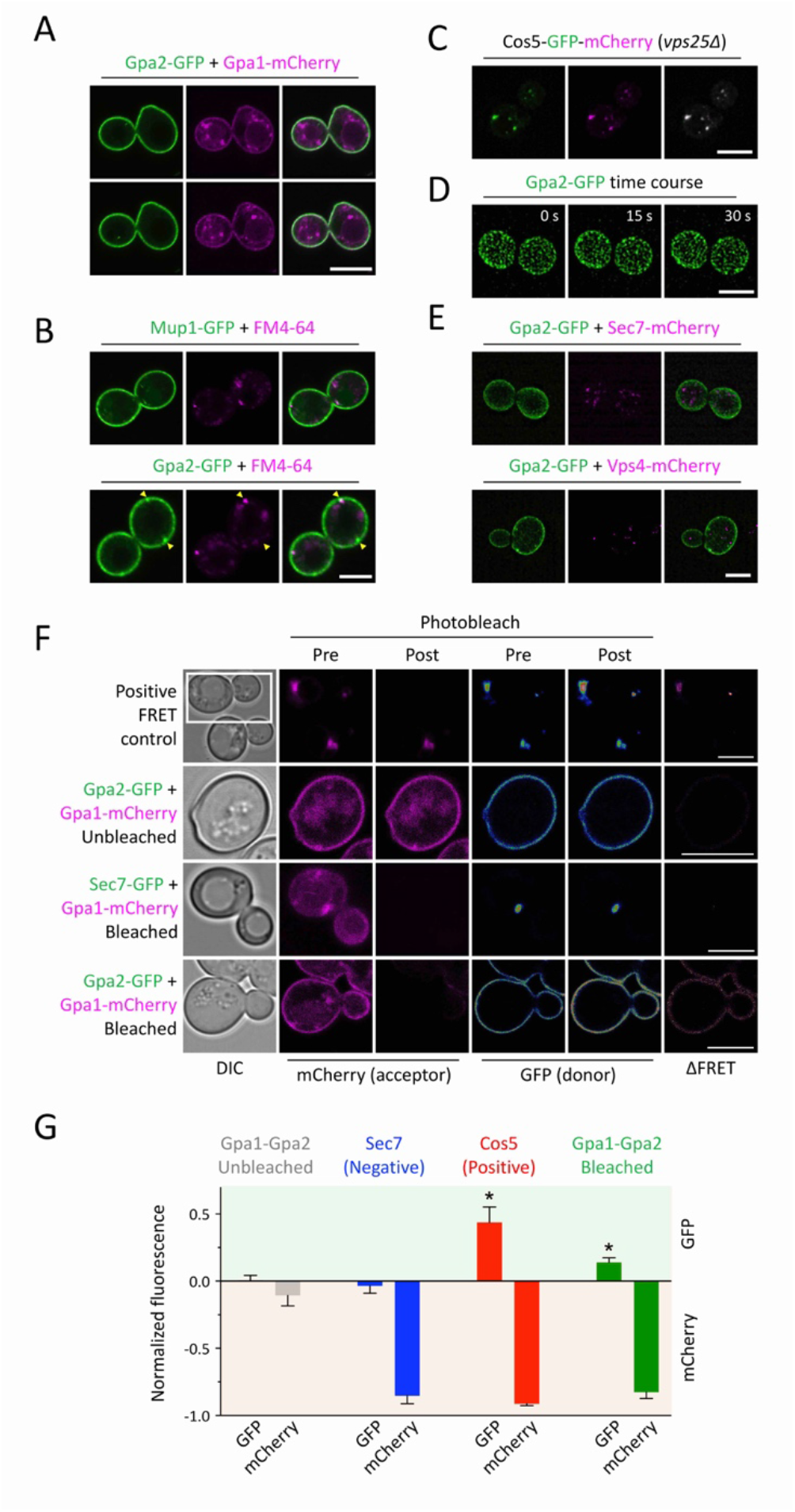
Gpa2 chiefly localises to the PM where it interacts with Gpa1. **A)** Wild-type cells co-expressing Gpa2-GFP and Gpa1-mCherry were imaged by confocal microscopy. **B)** Cells expressing either Mup1-GFP or Gpa2-GFP were labelled with 40 µM FM4-64 for 3 minutes followed by quenching of extracellular dye with 2.4 µM SCAS and confocal imaging **C)** 4D Apotome SIM experiments were performed for *vps25Δ* cells expressing a dual GFP-mCherry tagged Cos5. **D)** 3D time lapse Apotome SIM microscopy was used to image wild-type cells expressing Gpa2-GFP with 42 z-stack slices to cover fluorescence across depth of cells. Maximum projections of the top 10 z-slices are shown for top-focussed, surface labelled signal. **E)** Imaging was performed as described in (**D**) for wild-type cells co-expressing Gpa2-GFP with either Sec7-mCherry (upper) or Vps4-mCherry (lower), with maximum intensity projections generated across all 42 z-stack slices for each sample shown. **F)** Acceptor bleaching experiments using the 561nm laser at 100% were performed with expressed fluorescent proteins indicated. Conditions were optimised using a positive control: *vps25Δ* cells expressing Cos5-mCherry-GFP with a 7 amino acid linker between mCherry and GFP designed to give maximal FRET signal (upper). Gpa2-GFP and Gpa1-mCherry experiment was controlled by assessing fluorescence from an unbleached sample and from colocalised Gpa1-mCherry and Sec7-GFP. GFP fluorescence pre- and post-bleach are shown in RGB LUT format, The GFP fluorescence of pre-bleach was subtracted from post-bleach image (ΔFRET) and the Fire LUT applied. **G)** Normalised fluorescence for GFP and mCherry pre- and post-post-bleach datasets from (F) were quantified from n=3 experiments. Scale bar, 5 µm.

**Figure 11:**
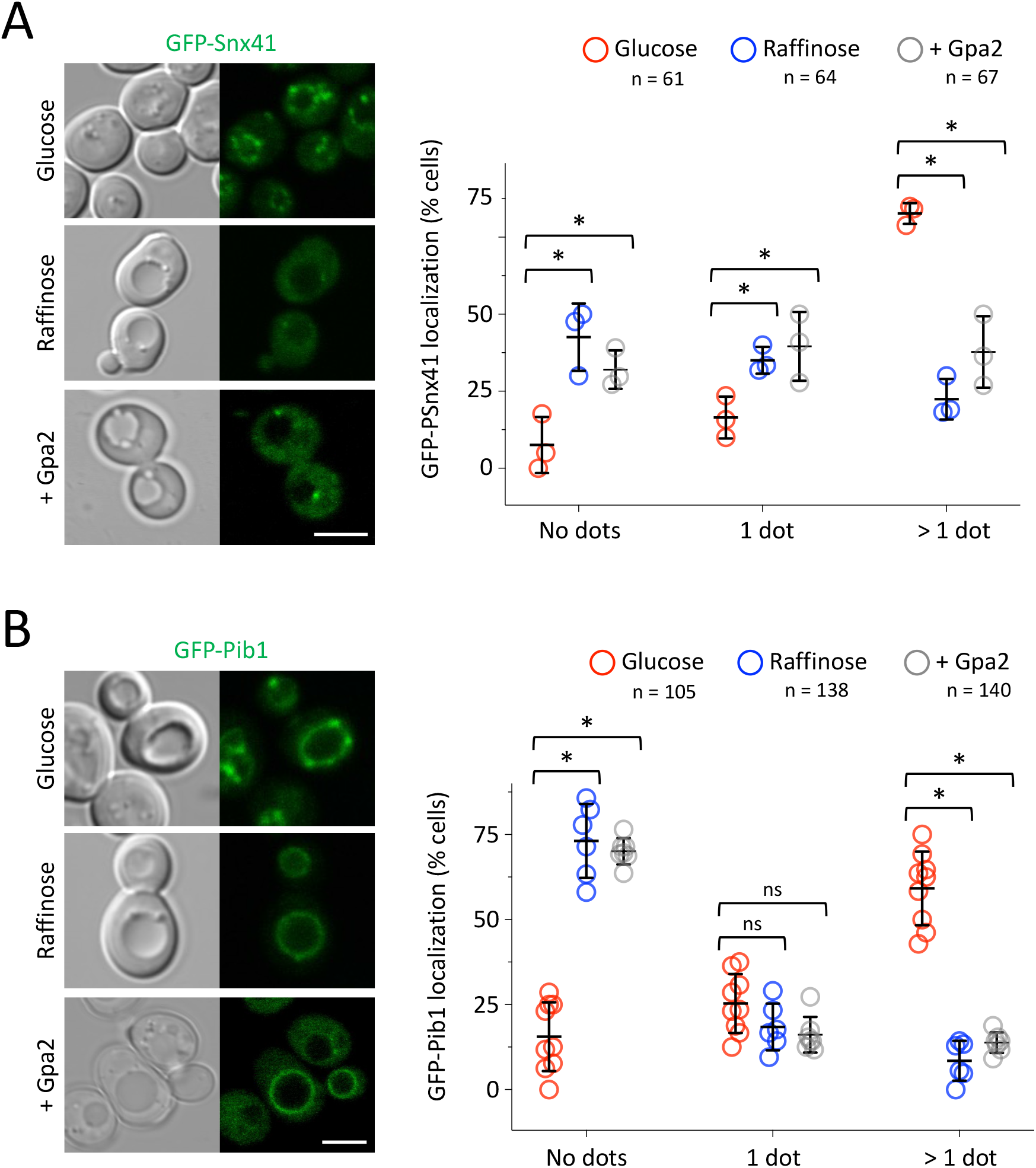
Glucose starvation and Gpa2 overexpression. **A-B)** Wild-type cells expressing GFP-Snx41 (A) and GFP-Pib1 (B) were imaged in glucose replete media, following 2 hours raffinose exchange, and in cells over-expressing Gpa2. Quantifications for each experiment were performed (right) by measuring how many dots of GFP-Snx41 were observed in each cell or by comparing the distribution of GFP-Pib1 dots as a percentage. Scale bar, 5 µm.

## DISCUSSION

Many protein and lipid trafficking itineraries are overhauled following acute nutrient depletion to equilibrate the energy balance of the cell and adjust metabolism appropriately for the change in extracellular conditions. Reduced recycling of material from endosomal compartments back to the PM benefits the cell by reducing anabolic load. Furthermore, surface proteins that are not recycled can be directed to the vacuolar degradation pathway instead and promote survival via catabolic processes. The division of labour between recycling and degradation pathways is not fully understood. Many surface proteins in yeast routinely accumulate in the vacuole, where fluorescent tags are stable, potentially overinflating the predominance of the well characterised degradation pathway. Ubiquitination of surface cargoes is sufficient to mediate their sorting via the ESCRT-driven multivesicular body pathway (Katzmann *et al*., 2001; Urbanowski and Piper, 2001; Babst *et al*., 2002b, 2002a). Most vacuolar cargoes are ubiquitinated by the Rsp5 E3-ligase and its cargo specific adaptors (O’Donnell and Schmidt, 2019; Sardana and Emr, 2021), but other ligases, such as Tul1 and Pib1, also contribute (Reggiori and Pelham, 2002; Li *et al*., 2015; MacDonald *et al*., 2017; Yang *et al*., 2020). Employing a ubiquitination reversal strategy, achieved by fusion of cargo, E3-ligases, or ESCRT-subunits to the catalytic domain of a deubiquitinating enzyme blocks cargo trafficking through the degradation pathway and allows focus on other endosomal trafficking events (Stringer and Piper, 2011; MacDonald *et al*., 2012b, 2015b). Here, we show that the GPCR Ste3, tagged with GFP and a DUb domain (Ste3-GFP-DUb) recycling is specifically inhibited following glucose starvation, with Ste3-GFP-DUb accumulating in endosomes. We assume efficient recycling of Ste3-GFP-DUb in wild-type cells maintains steady state signal at the PM, as increased internalisation through over-expression of the yeast AP180 adaptors (**Figure 1A**) or pheromone or temperature is insufficient to accumulate Ste3-GFP-DUb in endosomes (MacDonald and Piper, 2017). Additionally, irrespective of the internalisation rates of FM4-64 to endosomes (Vida and Emr, 1995), we find glucose starvation inhibits FM4-64 recycling back to the PM, measured as a percentage of internalised dye that subsequently effluxes (Wiederkehr *et al*., 2000). Therefore, we conclude that the Ste3-GFP-DUb reporter is a highly specific recycling reporter and that recycling of both protein and lipids is reduced in response to acute glucose starvation. A genetic screen for defects in Ste3-GFP-DUb recycling implicated the Gα subunit Gpa1 as required for recycling (MacDonald and Piper, 2017), which we confirmed by stably expressing Ste3-GFP-DUb in *gpa1Δ* cells, to show recycling is perturbed. Furthermore, *gpa1Δ* mutants cannot recycle FM4-64 as efficiently as wild-type cells and display either reduced PM signal or enhanced vacuolar sorting of various fluorescently tagged cargoes (**Figure 2**), the latter phenotype could be explained as an indirect consequence of reduced recycling.

Gpa1 has a role outside of surface GPCR signalling and can functionally connect with the yeast PI3K, comprised of Vps34 and Vps15, to simulate phosphatidylinositol 3-phosphate (PtdIns3P) synthesis at endosomes (Slessareva *et al*., 2006). Like *gpa1Δ* mutants, we find PI3K mutants are also defective in recycling (**Figure 4**). In addition to the role of PI3K in vacuolar trafficking of proteins through the biosynthetic (Robinson *et al*., 1988; Schu *et al*., 1993) and autophagy (Kihara *et al*., 2001; Wurmser and Emr, 2002) pathways, Vps34 generation of PtdIns3P is required for retrograde transport (Burda *et al*., 2002) and for the trafficking of proteins internalised from the PM and trafficked through the multivesicular body (MVB) pathway (Munn and Riezman, 1994; Katzmann, 2003). Our attempts to recover *vps15Δ* and *vps34Δ* cells from the Mat**α** collection, or to generate the mutants by homologous recombination, were unsuccessful. Localisation of the endogenously expressed Ste3-GFP-DUb reporter, which is mating-type specific, was therefore not possible. Instead, we performed experiments with Mat**a** *vps15Δ* and *vps34Δ* mutants, that are both extremely defective in FM4-64 recycling (**Figure 4A**), which one might reasonably expect following such dramatic abrogation of the endolysosomal system. However, our additional work suggest that the recycling pathway is modulated in response to more subtle PI3K regulatory effects. For example, we took advantage of an optimised hyperactive Vps34 allele (Vps34^EDC^) that stimulates over-production of PtdIns3P, which had previously been shown to upregulate retrograde trafficking, perturb late stages of autophagy and have no effect of MVB sorting (Steinfeld *et al*., 2021). Expression of hyperactive Vps34 resulted in defects in FM4-64 recycling (**Figure 4D**) suggesting, like the late stages of autophagy, recycling to the surface requires specific PtdIns3P regulation. This result also implies Gpa1-PI3K mediated recycling is distinct from retrograde trafficking routes via the Golgi (Ma *et al*., 2017; Best *et al*., 2020). The finding that a single point mutation in Vps15, which disrupts the interaction of the Gpa1 effector with PI3K (Heenan *et al*., 2009) was sufficient to disrupt recycling to a similar degree as *GPA1* deletion (**Figure 4F-H**) suggests that the recycling defects of *gpa1Δ* cells could be explained by improper PtdIns3P production. A potential connection between recycling defects observed during glucose starvation and PI3K-Gpa1 regulation was noted due to a constitutively active Gpa1^Q323L^ mutant also being defective in recycling (**Figure 5**). The screen that discovered signalling of Gpa1^Q323L^ requires PI3K also revealed *reg1Δ* cells, which lack the phosphatase subunit Reg1(Tu and Carlson, 1995; Sanz *et al*., 2000), gave the largest defect in signalling across all ∼5000 mutants tested (Slessareva *et al*., 2006). Although Gpa1 phosphorylation status is controlled by glucose associated enzymes (Elm1, Sak1, Tos3 and Glc7-Reg1), this has little impact on GDP or GTP binding (Clement *et al*., 2013), the latter being required for PI3K mediated production of PtdIns3P (Slessareva *et al*., 2006). Therefore, we hypothesised the reason *reg1Δ* cells suppress constitutively active Gpa1 was through a glucose sensitive transcriptional response mediated via Glc7 and the downstream transcriptional repressor Mig1 (DeVit and Johnston, 1999; Shashkova *et al*., 2017).

In support of this hypothesis, we observe recycling defects in both *reg1Δ* mutants (**Figure 5E, 5F**) but also in *mig1Δ mig2Δ* cells (**Figure 7C**) lacking downstream transcriptional repressor activity (Schüller, 2003). This suggested expression of an unknown inhibitor of Gpa1-mediated recycling would be de-repressed following glucose starvation or in *mig1Δ mig2Δ* mutants. The other yeast Gα subunit Gpa2 was the only candidate recycling inhibitor that both physically interacts with Gpa1 (Ho *et al*., 2002) but has also been proposed as a gene product repressed by Mig1 (Wollman *et al*., 2017). We confirmed *GPA2* meet these criteria by being transcriptionally upregulated ∼6.4 ± 0.7 fold following 1-hour raffinose exchange compared with glucose grown cells, and increasing ∼2.0 ± 0.3 fold in *mig1Δ mig2Δ* cells compared to wild-type (**Figure 7E**). We went on to show that Gpa2 over-expression is sufficient to reduce recycling efficiency of Ste3-GFP-DUb, FM4-64 and various fluorescently tagged surface cargoes (**Figure 9**). The finding that Ste3-GFP-DUb recycling defects in *mig1Δ mig2Δ* can be supressed by further deletion of *GPA2* supports the notion that Gpa2 is a recycling inhibitor controlled at the transcriptional level via Mig1 in response to glucose starvation (**Figure 7G**). The exact mechanisms of recycling inhibition via Gpa2 are not known, but we found little evidence of Gpa2 localising to endosomal structures that might suggest a direct role with Gpa1-PI3K (**Figure 10A - 10E**). Instead, based on the increased protein levels that concentrate at the surface during glucose starvation (**Figure 8**) and physical interaction of Gpa1 and Gpa2 (**Figure 10F, 10G**), we propose elevated levels of surface localised Gpa2 following glucose starvation divert Gpa1 from endosomes, thereby attenuating PI3K activity and recycling (**Figure 6B**). Although our steady state evidence is not sufficient to conclude that Gpa1 is sequestered by Gpa2 at the surface, we did find an increase of surface localised Gpa1 in the recycling mutant *rcy1Δ*, suggesting at least the distribution between surface and endosomal Gpa1 can be modulated in response to recycling efficiency (**Figure S3**). Alternatively, as Gpa1 is both palymitoylated and myristoylated (Song and Dohlman, 1996; Song *et al*., 1996), its ability to regulate endosomal lipids with PI3K may be required for its correct localisation. To test our model that elevated surface levels of Gpa2 inhibits recycling by reducing PI3K production of PtdIns3P, we examined the localisation of two proteins that bind endosomal membranes enriched in PtdIns3P. We found localisation of both the PX-domain protein Snx41 (Hettema *et al*., 2003) and the FYVE-domain protein Pib1 (Shin *et al*., 2001) were disrupted following glucose starvation, with marked reduction in membrane association (**Figure 11**). Support for our model that Gpa1 mediated PI3K activity is inhibited by the Gpa2 inhibitor comes from our discovery that simply over-expressing Gpa2 on a plasmid mimics glucose starvation and mis-localises Snx41 and Pib1.

We believe the mechanism described here serves as a medium-term transcriptional based solution in the initial hours of starvation. As surface proteome effects are also observed more rapidly, this response presumably integrates with faster acting regulation, potentially involving posttranslational modification of Rsp5 adaptors (Kahlhofer *et al*., 2021), and contributes to sustained accumulation in the vacuole over longer periods (Lang *et al*., 2014; Müller *et al*., 2015). Beyond the exact mechanism of Gpa2 inhibition, other important questions remain, such as the molecular function of Gpr1 and Ras2, which we confirmed are required for recycling (**Figure 2C**) whilst also being functionally associated with Gpa2 and glucose metabolism (Colombo *et al*., 1998, 2004). We speculate recycling inhibition via increased *GPA2* expression is distinct from its role with Gpr1-Ras to induce cAMP signalling (Xue *et al*., 1998; Kraakman *et al*., 1999), as over-expression of Gpa2 does not inhibit Gpr1 function, instead it triggers increased Ras signalling and cAMP accumulation (Nakafuku *et al*., 1988). Our model would suggest clathrin mediated endocytosis of surface cargoes is counter-balanced by efficient Gpa1-PI3K recycling in glucose replete conditions. In glucose, the tuneable inhibitor of recycling Gpa2 is transcriptionally repressed, collectively maintaining high levels of surface proteins at the PM for optimal growth (**Figure 6A**). Upon glucose depletion, the Mig1-dependent increase in endocytosis via yeast AP180 adaptors (Laidlaw *et al*., 2020), would complement decreased recycling via the Glc7-Reg1 > Mig1 > *GPA2* pathway to modulate the surface proteome to suit nutritional availability, increase lysosomal / vacuolar degradation and calibrate metabolic processes for cellular survival. G-protein regulators and PI3K orthologues are evolutionarily conserved (Pierce *et al*., 2002; Engelman *et al*., 2006), and although G-protein signalling is much more complex in animal cells, Gα_s_ have been shown to regulate endosomal trafficking and surface protein function (Colombo *et al*., 1992, 1994; Beron *et al*., 1995; Zheng *et al*., 2004; Beas *et al*., 2012). Therefore, this mode of surface protein regulation in response to nutrition may be conserved in mammalian cells.

## METHODS

### Reagents

Supplemental tables are included to document use of yeast strains (**Table S1**), plasmids (**Table S2**) and statistical tests (**Table S3**).

### Cell culture

Yeast cells were cultured in either rich media (yeast extract peptone dextrose (YPD); 2% glucose, 2% peptone, 1% yeast extract) or synthetic complete minimal medium (SC; 2% glucose, yeast nitrogen base supplemented with amino acid / base dropout mixtures. Cultures were routinely prepared in serial dilution overnight so that cells were harvested for downstream experiments from early / mid-log phase log phase (OD_600_ = <1.0). For glucose starvation experiments, 2% glucose media was washed 3x and exchanged with either identical media lacking any carbon source (no sugar) or media of the same receipe but instead containing 2% raffinose instead of glucose. KanMX and ClonNAT strain selections were performed in rich media containing 250 μg/ml geneticin/G418 (Formedium) or 150 μg/ml Nourseothricin (Jena Biosceince), respectively. GFP and mCherry fusions of *SEC7* were created with a methotrexate cassette selected on 20 mM methotrexate (Alfa Aesar) supplemented with 5 mg/ml sulphanilamide. The *loxP* flanked cassette was then excised by *TEF1*-Cre expression and plasmid removal, as described (MacDonald and Piper, 2015). Expression of plasmids from the *CUP1* promoter in appropriate selective media was induced by the addition of Copper chloride (typically 20 - 100 µM).

### Confocal microscopy and Förster resonance energy transfer (FRET)

Cells were typically harvested from mid-log phase SC minimal media cultures and prepared for confocal microscopy on Zeiss laser scanning confocal instruments (LSM710 or LSM880 equipped with an Airyscan) using a Plan-Apochromat 63x/1.4 Differential Interference Contrast (DIC) objective lens. The fluorescent proteins GFP, mGFP and mNeonGreen were excited using the 488nm line from an Argon laser their emission collected from 495 – 550 nm. Fluorescent protein mCherry and dye FM4-64 were excited using 561nm line from a yellow DPSS laser and the emission collected 570 – 620 nm. Acceptor bleaching was used to determine the occurrence of FRET. The targeted bleaching of the mCherry (acceptor) using 561nm laser was used to dequench any FRET GFP (donor). Bleach acquisition was controlled by Zeiss Zen FRAP module where a short time-lapse series was taken before and after the bleaching of mCherry. The change in GFP intensity was measured and the data exported for visual representation in Fiji or plotted as graphs in GraphPad Prism (v9.0.2).

### Apotome Structured Illumination Microscopy

Cells were imaged on a Zeiss Elyra 7 system using Plan-Apochromat 40x/1.4 oil objective lens. Multi-coloured acquisition was performed sequentially to minimise cross talk between channels. The fluorescent images were capture on 2 PCO Edge sCMOS cameras attached to a DuoLink motorised dual camera adapter and the colour split using the secondary beam splitter BP420-480 + BP470-640 + LP740. The fluorescent protein mCherry was excited using 561nm laser line and emission collected from 570 to 640nm. The fluorescent protein GFP was excited using 488nm laser line and emission collected from 490 to 570nm. Apotome acquisition was set to collect 5 phase images with 25ms camera exposure time. Yeast cells were optical Z sections using step size optimised for “Leap” acquisition. For time lapse experiments z stacks were collected with an interval of 4.3 seconds. Apotome phase images were processed using Zeiss Zen Black software set to 3D SIM Leap. The alignment between the colour channels was further improved by taking a Z stack of multi-colour TetraSpec microspheres (ThermoFisher) that was used to generate an alignment matrix using the “Channel alignment” tool in Zen Black and applied to the time lapse data.

### Image analysis

Micrographs were processed by Zeiss Zen and Fiji software. Images were processed using Zen software (Zeiss) and were further modified (e.g. coloured, merged) using Fiji.

### Halo mating assay

Mat**a** wild-type cells and *gpa1Δ* mutants transformed with either empty vector or Gpa1-mCherry cells were grown to saturation overnight, diluted in fresh SC media and grown for ∼6 hours. Equal amounts of cells were estimated by measuring the OD_600_ of the culture, harvested and resuspended in 50 µl sterile water before spotting onto lawns of Mat**α** *bar1-1* mutant cells. Lawns were created from mid-log phase cultures spread on YPD solid agar and left to dry for several hours. Plates were incubated at 30 degrees for 2 days before the area of growth inhibition was measured using ImageJ (NIH) for each spot of Mat**a** cells. These area measurements were normalised to wild-type cells from the same plate and the average plotted for 4 biological replicates.

### Yeast RNA extraction

For gene expression analysis following glucose starvation wild-type cells were grown to mid-log phase in YPD before being split and incubated in 10 ml YPD (dextrose) or 10 ml YPR (raffinose) media for 1 hour prior to harvesting. For experiments to test the role of Mig1 / Mig2, wild-type and *mig1Δ mig2Δ* cells were grown to mid-log phase in 10 ml YPD before harvesting. Spheroplasting of harvested yeast cells was performed for 2 minutes in lysis buffer (1 M sorbitol 100 mM EDTA, 0.1% β-mercaptoethanol) containing 25 units of zymolyase (Zymo Research). RNA extraction was performed with an RNeasy kit (QIAGEN) including additional DNaseI treatment using a TURBO DNA-free kit (Invitrogen).

### Quantitative reverse transcription PCR (RT-qPCR)

cDNA was synthesised from 5μg extracted RNA with SuperScript IV reverse transcriptase (Invitrogen) using 50ng/μ random hexamers and 10mM dNTPs. 5-minute incubations at 65°C were carried out before 100mM DTT and Ribonuclease inhibitor added and the Superscript IV reverse transcriptase to initiate the qPCR reaction (10 mins 23°C; 10 mins 55°C; 10 mins 80°C) immediately. In order to amplify *GPA1*, oligonucleotides 361 (5’ ACATCGGCTCGTCCAAATTC) and 362 (5’ TCTGGTCGTATTCACTCATTGC) were used. To amplify *GPA2* oligonucleotides 365 (5’ CAATGGGCCTAACGCATCG) and 366 (5’ GGGTCTGTAATTGGGCGAAG) were used. All experiments were compared to *ACT1* reference gene amplified by oligonucleotides 207 (5’ CTCCACCACTGCTGAAAGAG) and 208 (5’ GCAGCGGTTTGCATTTCTTG). Single product amplification was confirmed by PCR using genomic DNA as a template, and near-100% amplification efficiencies were confirmed (102.2±0.12% for *GPA1*, 101.5±0.03% for *GPA2*, and 100.6±0.08% for *ACT1*) by duplicate qPCR reactions on a standard curve of known input quantity. qPCR reactions were performed on 20 ng cDNA, including relevant negative controls, in 20 µl reactions containing 350 nM of each primer, and 10 µl Fast SYBR(tm) Green Master Mix (ThermoFisher Scientific). The QuantStudio 3 system (Thermofisher) was used for reactions under the following conditions: 40 cycles of 95°C for 1 second, 20 seconds 60°C, before a continuous ramp from 60°C to 95°C at a rate of 0.1 °C/S for melt curve analysis. Expression of *GPA1* and *GPA2* under indicated conditions were quantified using the comparative Ct (ΔΔCt) method, relative to the expression of the housekeeping gene *ACT1* and normalised to control sample (glucose for raffinose comparisons and wild-type cells for comparison with *mig1Δ mig2Δ* mutants).

### Yeast genomic DNA extractions

Yeast for genotyping and genome sequencing was grown to mid-log phase in YPD, before being harvested and resuspended in 50 mM Tris.HCl 20 mM Ethylenediaminetetraacetic acid (EDTA) and then 3 µl β-mercaptoethanol, 10 µl zymolyase and 1 mg/ul RNase (QIAGEN). This was incubated in a 37 °C shaker for 1 hour before the addition of Proteinase K (10 µl, QIAGEN, >600 mAU / ml) and left at 55 °C for 1 hour. 500 µl of Phenol:Chloroform:Isoamyl alcohol was added and the solution was vortexed for 5 minutes before being spun at 15 000 rpm for 5 minutes. The aqueous layer was transferred to a fresh Eppendorf and this was repeated two more times. The final aqueous layer was transferred to a fresh Eppendorf and 50 µl 3M Sodium Acetate was added. DNA was precipitated by the addition of 1 ml 100% ethanol and spinning (10 °C 15000rpm) for 10 minutes. The pellet was then washed with 70% ethanol before residual ethanol was removed. The pellet was then dissolved in 100 µl TE Lite and left to resuspend overnight at room temperature.

### Yeast genome sequencing

Prior to sequencing library generation, genomic DNA samples extracted as above were subject to an additional clean up step, by binding samples with a 1.5 X volume of AMPure XP beads (Beckman Coulter), washing twice with 70 % ethanol, and eluting into fresh TE. Libraries were then prepared from 500 ng genomic DNA using the NEBNext Ultra II FS DNA library prep kit for Illumina (New England Biolabs), according to the manufacturer’s instructions, and using a 14 minute, 37 °C incubation for DNA fragmentation, and 3 cycles of PCR for incorporation of unique dual indices (NEBNext multiplex oligos for Illumina) to the final libraries. Following library quantitation and quality assessment using the Agilent Tapestation, libraries were pooled at equimolar ratios, and subject to 150 base paired end sequencing on an Illumina NovaSeq at Novogene, Europe.

### Genome Alignment and Variant Detection

Illumina paired end reads were quality trimmed and verified using Cutadapt v3.4 (Martin, 2011) and FastQC (Andrews, 1010) respectively, before alignment to the *Saccharomyces cerevisiae* reference strain S288C using BWAmem (Li, 2013). Any mistakes in mate pairing and duplicates were detected and removed, and read sorting was performed using samtools v1.11 (Danecek *et al*., 2021). Resultant bam files were visually inspected in IGV viewer (Robinson *et al*., 2011) <u>for verification of the modifications to the *GPA1* locus (*gpa1Δ::kan*^*r*^</u>) in both the BY4741 Mat**a** and BY4742 Mat*α* <u>strain backgrounds</u>. The three bam files were arranged into genomic positions using samtools and variants were identified using VarScan v2.9.3 (Koboldt *et al*., 2009). Variants were analysed with SnpEff (Cingolani *et al*., 2012) for annotation of variants as well as Variant Effect Predictor (McLaren *et al*., 2016) for confirmation.

### Flow cytometry

Fluorescence intensity of different strains stably expressing Ste3-GFP-DUb was measured using a CytoFLEX flow cytometer (Beckman Coulter) with 488 nm laser excitation, 525 / 40 nm emission filter and avalanche photo diode detector. Approximately 75,000 cells in culture medium per sample were analysed with logical gating (forward/side scatter: single cells) used to quantify GFP fluorescence-positive yeast cells. Sample analysis was performed using the software FCS Express v7.04 (DeNovo software).

### FM4-64 recycling assay

Cells were grown to mid-log phase in YPD, or SC-Ura minimal media when plasmid selection was necessary, before 1ml of cells (OD = 1.0) was harvested and incubated in 100 µl YPD containing 40 µM FM4-64 dye (*N*-(3-Triethylammoniumpropyl)-4-(6-(4-(Diethylamino) Phenyl) Hexatrienyl) Pyridinium Dibromide) dye for 8 minutes at room temperature. Cells were then washed 3x in cold SC media, with each wash left for 3 minutes on ice, before final wash was concentrated in 100 µl SC media in preparation for flow cytometry. For raffinose and glucose starvation media, the same rich and SC media was used with glucose exchanged with either no sugar or 2% raffinose. 20 µl of concentrated cells were brought up in 3 mls room temperature SC medium, and approximately analysed by flow cytometry at 1000 – 2500 cells per second using an LSR Fortessa X20 instrument (BD Biosciences). FM4-64 intensity was measured over a 10-minute period with 561nm laser excitation and emission filter 710 / 50. Measurements from 488 nm laser excitation with 530 / 50 nm emission filter were also recorded for monitoring background autofluorescence. Any comparisons are performed from cells labelled at the same time in the same media, with empty vector controls included when effects from plasmids assessed.

### Image quantification for Mig1-GFP

The nuclear signal of Mig1-GFP was calculated as a percentage of nuclear / total signal in the green channel, as shown in (**Figure S2**). Briefly, whole cells were segmented based on DIC image using the Cell Magic Wand Plugin (Min = 8, Max = 300, roughness = 2.0) and used to calculate the total (nuclear plus cytoplasmic) signal for each cell in the Mig1-GFP / Green channel. The Nrd1-mCherry signal was used with otsu threshold to segment the nucleus based on the red channel and these regions of interest were applied to measure nuclear Mig1-GFP from the green channel.

### Statistical analyses

Unpaired Student’s *t*-tests were performed using GraphPad Prism v8.3.1. to compare the statistical significance between experimental conditions, with an asterisk (*) used to denote p-values of <0.05 or less, as mentioned in specific figure legends, or (ns) used to define differences that are not significant.

## Supporting information

Table S1: Yeast Strains used in this study

Table S2: Plasmids used in this study

Table S3 - Statistical analysis

## ACKNOWLEDGMENTS

We would like to thank staff at the York Bioscience Technology Facility for technical assistance. We are grateful to Lois Weismann and Noah Steinfeld (University of Michigan) for reagents to manipulate yeast PI3-kinase activity and Chris Stefan (UCL) for fruitful discussions. This research was supported by a Sir Henry Dale Research Fellowship from the Wellcome Trust and the Royal Society 204636/Z/16/Z (CM).

