## Supplemental Figures and Legends for "Endosomal cargo recycling mediated by Gpa1 and Phosphatidylinositol-3-Kinase is inhibited by glucose starvation"

- **Figure S1:** Fluorescently tagged Gpa1 complements the mating defect of gpa1Δ mutants
- **Figure S2:** Segmentation and quantification of nuclear Mig1-GFP
- **Figure S3:** Gpa1 and Gpa2 colocalisation in wild-type and recycling mutant cells
- **Figure S4:** Fluorescently tagged Gpa2 complements the small cell size defect of *gpa2Δ* mutants
- **Figure S5:** Flow cytometry analysis focussed specifically on transformed cells
- **Figure S6:** Localisation of Cos5-GFP in wild-type and MVB sorting mutants
- **Figure S7:** Apotome SIM localisation experiments
- **Figure S8:** FRET measurements to document surface interaction between Gpa1 and Gpa2

### SUPPLEMENTAL REFERENCES

### SUPPLEMENTAL TABLES

- **Table S1:** Yeast strains used in this study
- **Table S2:** Plasmids used in this study
- **Table S3:** Results from statisitical tests

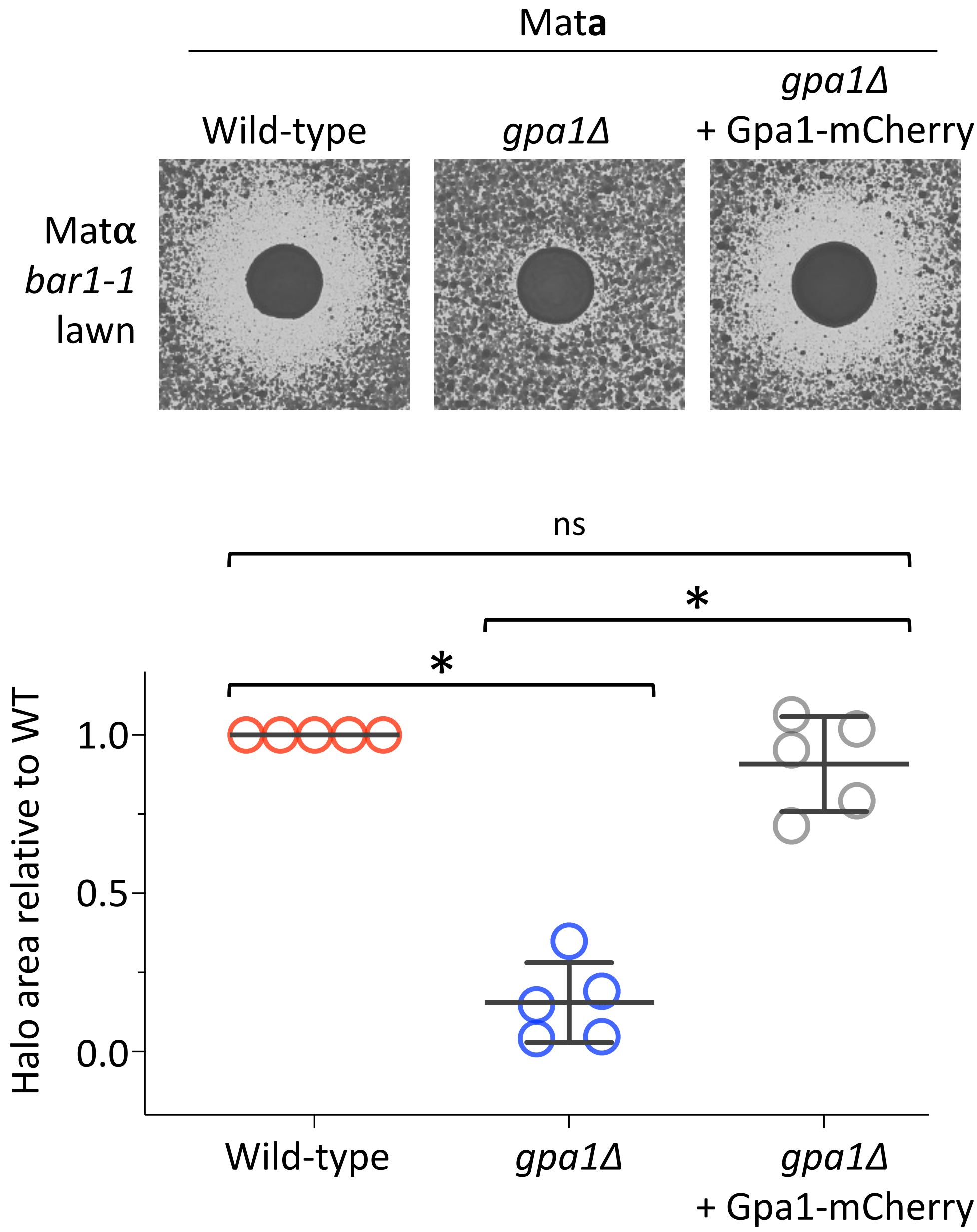


**Figure S1: Fluorescently tagged Gpa1 complements the mating defect of *gpa1*Δ mutants**
Matα cells harbouring the *bar1-1* allele were grown to mid-log phase then plated as a lawn on rich media agar plates and left to dry. Mat**a** cells grown to exponential phase were spotted onto the dried lawns and grown for 2 days (upper). The 'halo' area of growth inhibition was measured using imageJ and normalised to a wild-type control on the same plate (lower). n = 5, Student's *t*-test was used to determine significance (* denotes $p < 0.0001$).

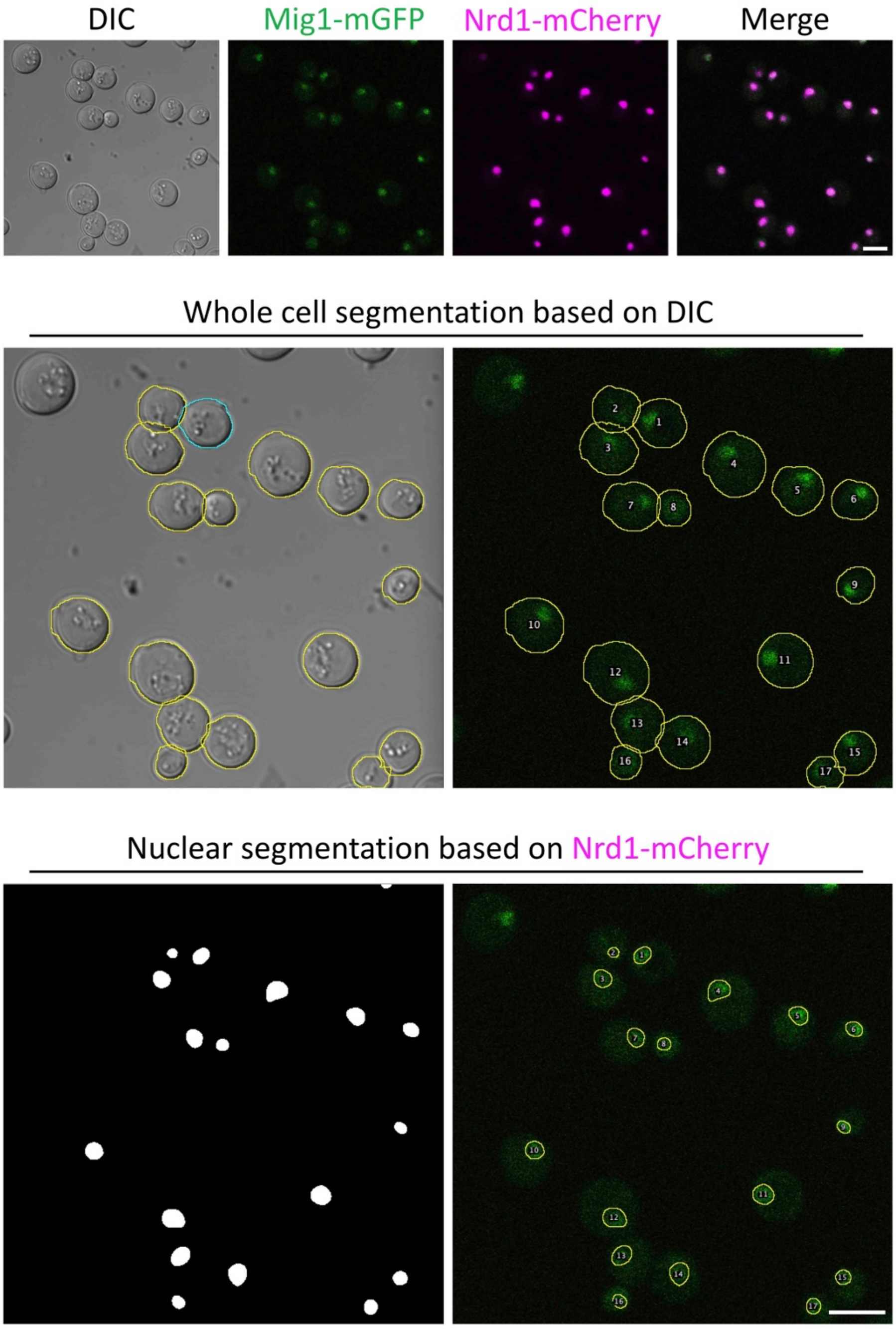


**Figure S2: Segmentation and quantification of nuclear Mig1-GFP**
Wild-type cells stably expressing Mig1-mGFP and Nrd1-mCherry were grown to mid-log phase in SC media containing 2% glucose (upper). Using the Cell Magic Wand Plugin (Fiji) set to roughness = 2.0 whole cells were identified from the DIC image, and these segmented regions of interest (ROIs) were applied to the green channel image to calculate the overall Mig1-mGFP fluorescence (middle). For nuclear specific localisations, Ostu segmentation was applied to the red Nrd1-mCherry channel to create nuclear ROIs that were then applied to the same Mig1-mGFP fluorescence cannel. Percentage Mig1-mGFP nuclear / total fluorescence was calculated for individual cells across multiple imaging experiments (n = 3). The same process was also performed for cells grown in SC media containing raffinose instead of glucose. Scale bar, 5 μM.

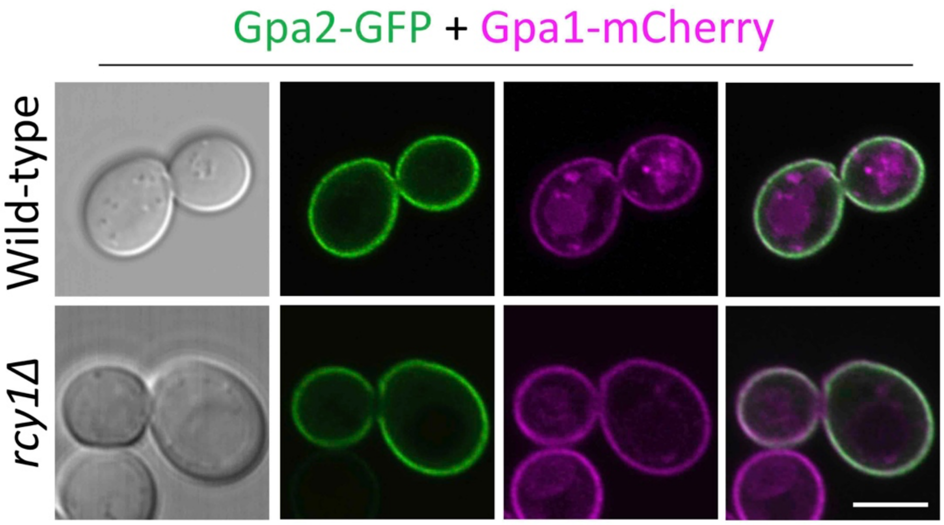


**Figure S3: Gpa1 and Gpa2 colocalisation in wild-type and recycling mutant cells**
Wild-type and *rcy1*Δ cells co-expressing Gpa2-GFP and Gpa1-mCherry expressed from the *CUP1* promoter induced by addition of 50 μM copper chloride to the media were imaged using confocal microscopy. Scale bar, 5 μm.

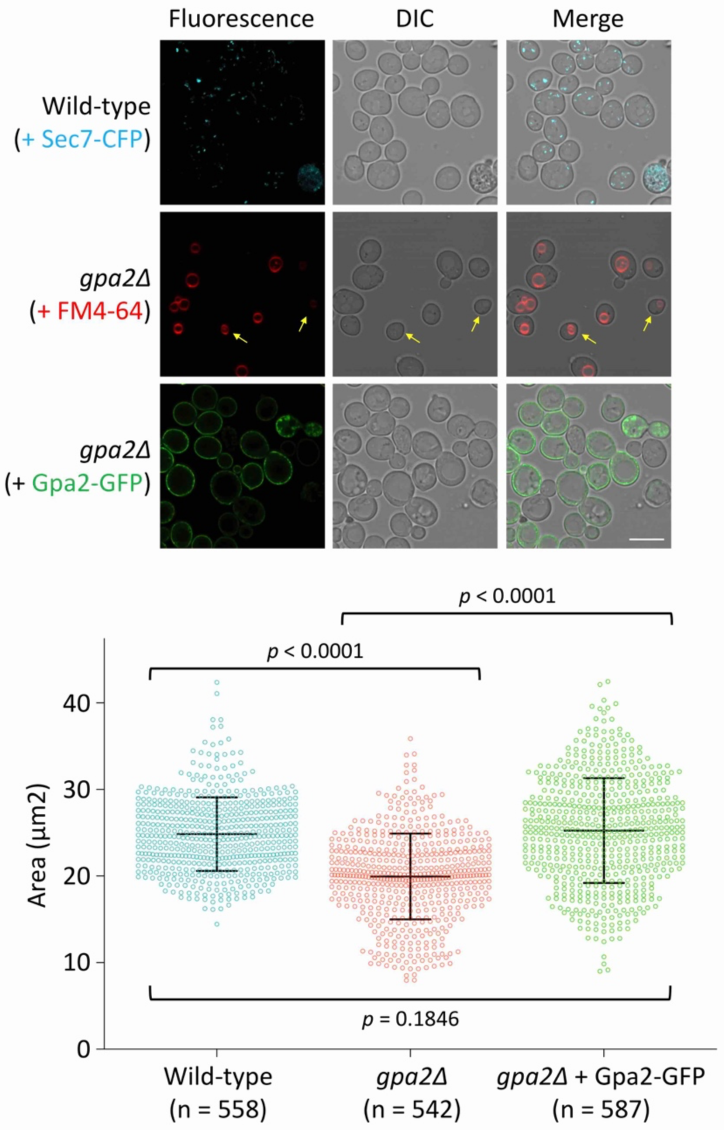


**Figure S4: Fluorescently tagged Gpa2 complements the small cell size defect of *gpa2*∆ mutants**
Wild-type indicated cells were grown to mid-log phase and prepared for fluorescence microscopy. To distinguish cell populations, wild-type cells expressed Sec7-CFP, the vacuoles of *gpa2*∆ nulls transformed with a vector were labelled with FM4-64 for 1 hour followed by a 30 minute dye-free chase period, and *gpa2*∆ nulls were expressing Gpa2-GFP (upper). Following segmentation using Cell Magic Wand tool in ImageJ, cellular area was measured and plotted (lower). As the cell size varies throughout cell cycle, a large number of cells (n > 500) were quantified for each condition. Scale bar, 5 µm. *p* value from Student's *t*-test comparisons indicated.

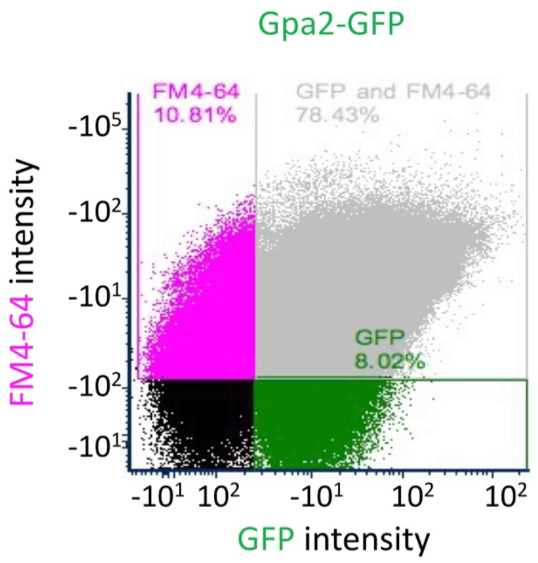


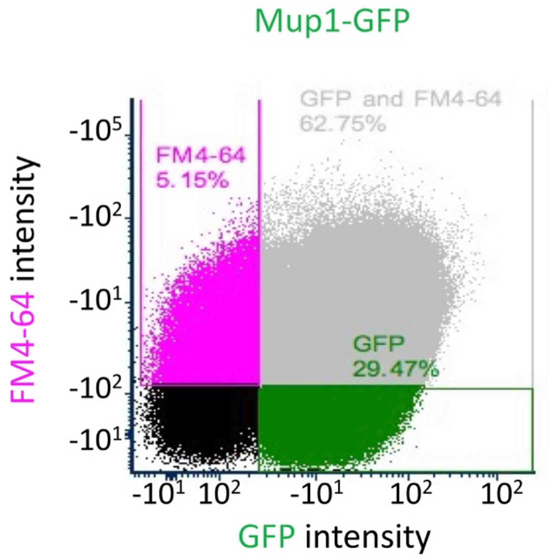


**Figure S5: Flow cytometry analysis focussed specifically on transformed cells**
Wild-type cells expressing either Gpa2-GFP (lower) or Mup1-GFP (lower) were grown to mid-log phase before preparation for FM4-64 efflux assays (see methods). Briefly, cells were loaded with FM4-64 dye before excess dye was washed with ice cold media. Flow cytometry measurements of cells upon a return to room temperature media was recorded and gates set to only calculate FM4-64 fluorescence from cells also co-expressing either Gpa2-GFP or Mup1-GFP. A decrease in fluorescence is plotted in Figure 9C, calculated from the mean fluorescence from the first 10 seconds of recording, considered 100%, and then applied to all subsequent measurements over the 10- minute period of continuous flow / measurements.

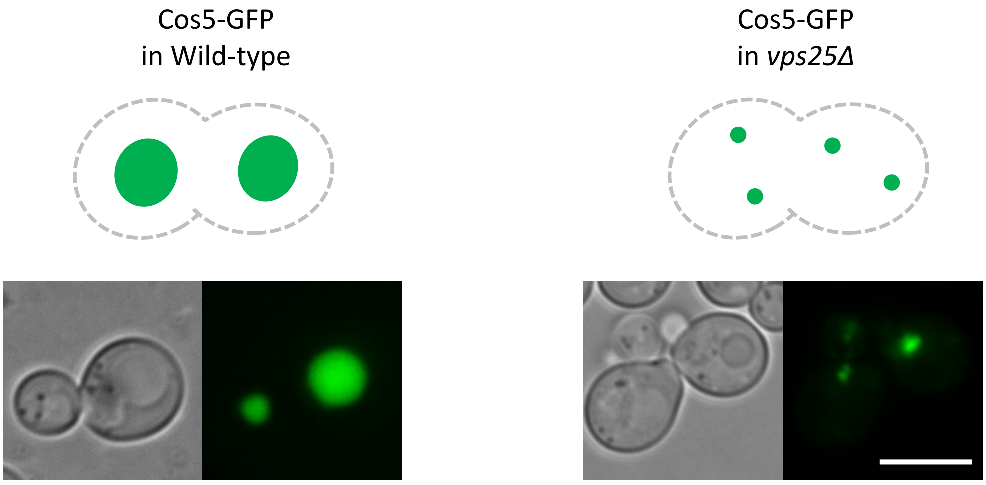


**Figure S6: Localisation of Cos5-GFP in wild-type and MVB sorting mutants**
Cos5-GFP was expressed from the *TDH3* promoter in either wild-type cells (left) or in *vps25*Δ mutants, that are defective in multivesicular body sorting and accumulate cargoes in aberrant endosomes (right). Scale bar, 5 μM.

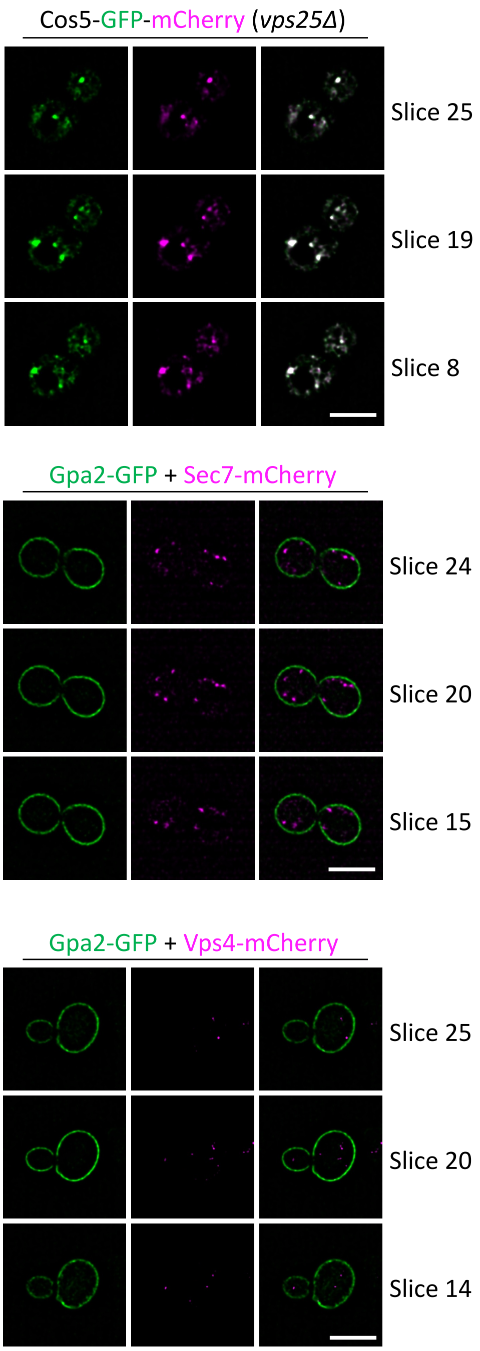


**Figure S7: Apotome SIM localisation experiments**
4D Apotome SIM was achieved across 42 z-stacks (distance 0.126µm) repeated over 100 time slices, each of 4.3 seconds with no interval period. This approach was used to image: a dual tagged version of Cos5, carrying both GFP and mCherry at the C-terminus was expressed in *vps25*Δ (upper), and wild-type cells co-expressing Gpa2-GFP with either Sec7-mCherry (middle) or Vps4-mCherry (lower). Scale bar, 5 µM.

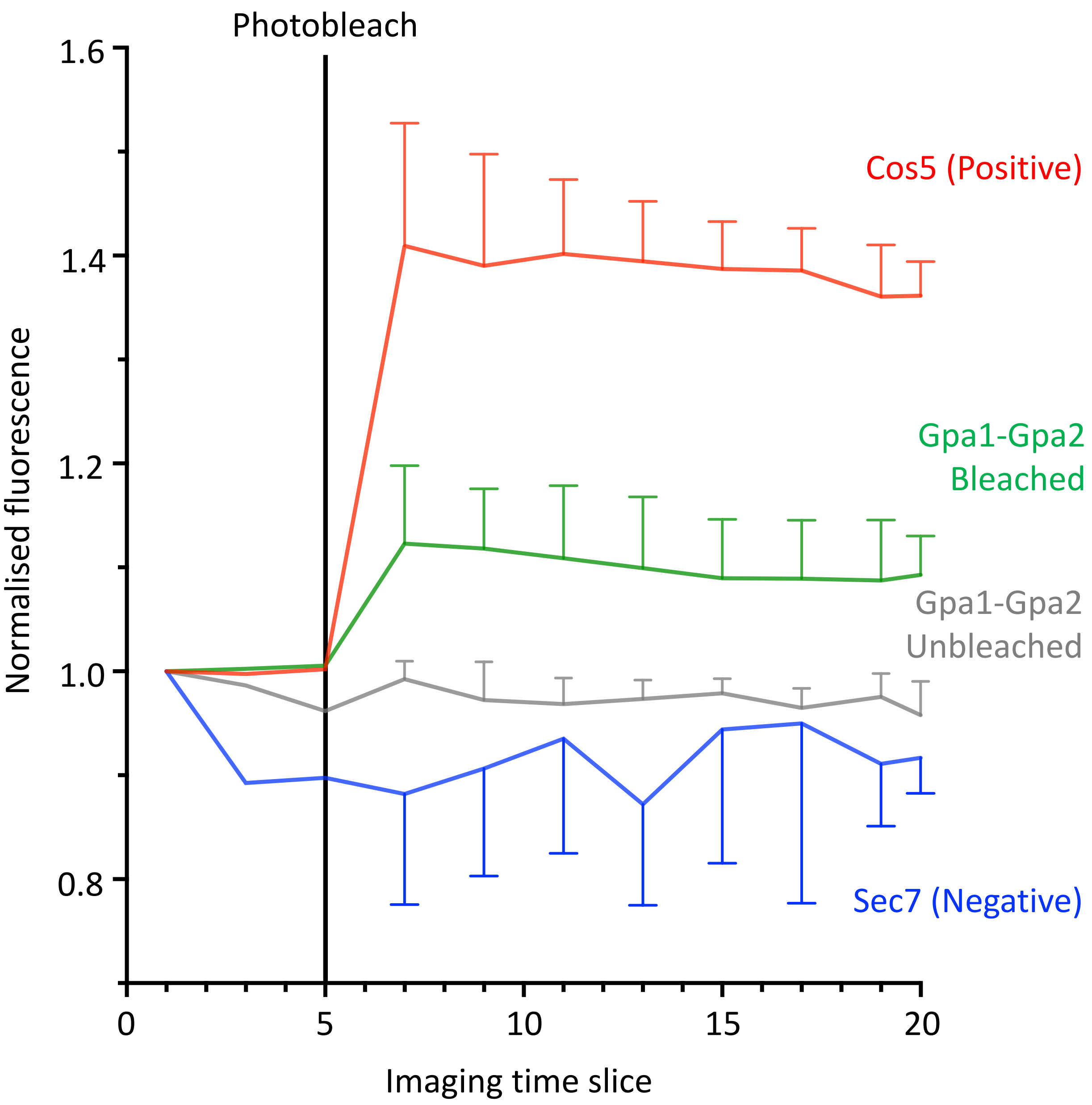


**Figure S8: FRET measurements to document surface interaction between Gpa1 and Gpa2**
Acceptor photobleaching experiments were performed as described in methods and shown in Figure 10F, 10G. To ensure stable measurements were observed, before and after bleaching was recorded in both red and green channels. The average change in GFP fluorescence is shown with error bars representing standard deviation from 3 replicates. As a positive control, Cos5 labelled with mCherry and GFP on the same molecule separated by a 7 residue linker was imaged (red). Gpa1-Cherry bleaching showed a positive FRET signal when colocalised with Gpa2-GFP (green) but not when colocalised with the *trans*-Golgi marker Sec7-GFP (blue). As a further control, Gpa2-GFP did not show any FRET signal in cells co-expressing Gpa1-mCherry but that were not subjected to bleaching.
